# Improved Anisotropic Network Models for Membrane Protein Dynamics and Mechanosensitive Ion Channels

**DOI:** 10.1101/2025.05.22.654704

**Authors:** Zhongjie Han, Lei Wang, Chen Song

## Abstract

Mechanosensitive (MS) ion channels are crucial for translating mechanical stimuli into cellular responses; however, their dynamic properties pose significant challenges for experimental investigations. This study introduces two enhanced Anisotropic Network Models (ANMs) that incorporate Membrane Contact Probability (MCP) to more effectively capture the influence of the cell membrane on protein dynamics. MCP-ANM and MCP-mANM (multiscale ANM) outperform traditional ANM models that do not explicitly consider membranes when predicting the flexibility of various types of membrane proteins. Furthermore, by integrating MCP-mANM with Perturbation Response Scanning (PRS), we are able to efficiently simulate the conformational changes of MS proteins subjected to different mechanical forces. This methodology allows for the examination of various gating mechanisms, including force-from-lipids and force-from-tether, across multiple MS ion channels such as NOMPC, MscS, and PIEZO. The calculated mechanosensitivity of the MS channels aligns well with the experimental findings. These results demonstrate the effectiveness of MCP-incorporated ANM in linking mechanical stimuli to protein dynamics, providing a reliable and efficient framework for understanding the mechanotransduction and gating mechanisms of MS ion channels.

## Introduction

Mechanosensitive (MS) ion channels are a specialized class of membrane proteins that respond to mechanical stimuli, including membrane tension, cell compression, and pressure. These stimuli facilitate a variety of physiological processes, such as touch sensation, hearing, and the regulation of cellular volume. ^1,2^ These MS channels, including well-known examples such as Mechanosensitive Channel of Small Conductance (MscS), TWIK-related K^+^ channel (TREK), Piezo-type mechanosensitive ion channel (PIEZO), and Hyperosmolality-gated calcium-permeable channel (OSCA), undergo conformational changes under mechanical stimuli to regulate ion flux, thereby maintaining cellular homeostasis under external stress. ^2^ Dysfunction of MS channels is implicated in various diseases, including hypertension, osteoporosis, and neuropathic pain, highlighting their importance as therapeutic targets. ^3–5^

Despite their significance, the study of MS ion channels remains challenging due to their dynamic and force-sensitive nature. Techniques such as patch-clamp electrophysiology provide detailed insights into ionic currents; however, they encounter difficulties in accurately correlating mechanical forces with gating behaviors. ^6^ Structural biology methods, including X-ray crystallography and cryo-electron microscopy (cryo-EM), provide information on static conformational states but often fail to capture the dynamic transitions that are essential for mechanosensation. ^7^

Computational methods such as molecular dynamics (MD) simulations^8^ have been widely used as a powerful tool to explore the dynamics of the MS channel at the atomic level, revealing insights into conformational changes and activation mechanisms. For example, Reddy et al. employed MD simulations to explore the gating mechanism of MscS channel, illustrating how membrane tension triggers structural rearrangements that open the pore. ^9^ More recently, Britt et al. have shown that MscS is modulated by lipid interactions, emphasizing the importance of membrane composition in channel behavior. ^10^ Similarly, Rogacheva et al. used unbiased MD simulations to uncover an asymmetric opening mechanism in the MscL channel under tension, which involves sequential conformational shifts. ^11^ Clausen et al. applied atomistic MD simulations to show that membrane-mediated structural changes couple the proximal C-terminus to the selectivity filter in TREK-2, thereby regulating its mechanosensitive gating.^12^ For the OSCA channel, Zhang et al. resolved its high-resolution structure using cryo-EM and used MD simulations to gain initial insights into its gating behavior. ^13^ Building on this, Han et al. integrated cryo-EM with MD simulations to reveal that mechanical stress induces the formation of a lipid-lined pore, emphasizing the role of lipid–protein interactions. ^14^ For the No Mechanoreceptor Potential C (NOMPC) channel, Wang and Duan employed MD simulations to show that compression and rotation of the anchor domain drive a unique ‘push-to-open’ and ‘twist-to-open’ gating mechanism. ^15,16^ However, MD simulations remain computationally intensive, particularly for large MS proteins such as PIEZO, which necessitate extended simulations to fully observe gating events. ^17–19^ Consequently, MD simulations are impractical for high-throughput analyses across a diverse range of MS proteins.

Coarse-grained models, such as elastic network models (ENMs), offer a computationally feasible alternative by focusing on large-scale functional conformational changes without the detailed resolution of atomistic MD. ^20^ Such methods can successfully capture low-frequency motions that characterize large conformational shifts in proteins. ^21^ However, despite the advantages of traditional Elastic Network Model (ENM) approaches, they encounter limitations in incorporating environmental factors critical for protein function, particularly in the context of membrane environments for membrane proteins. Previous adaptations of the Anisotropic Network Model (ANM, an extension of ENM that considers motion direction^22^), such as the implicit membrane Anisotropic Network Model (imANM), ^23^ attempt to account for the influence of the membrane by adjusting the spring force constants in ANM according to the positioning of the membrane. These adaptions have indeed improved the computational analysis of membrane proteins to a certain extent.^23^ Although these methods typically assign stiffer spring constants in the XY-plane (membrane plane) compared to the Z-direction to simulate membrane confinement, this simplified approach may not accurately reflect the full complexity of biological membranes, where the stiffness in the Z-direction could be greater, or the overall situation may be more intricate due to the complex interactions between proteins and membranes.

To address these limitations, we propose membrane contact probability (MCP)-based ANM models, MCP-ANM and MCP-mANM (multiscale ANM). These models incorporate the predicted membrane contact probability (MCP), as illustrated in Figure 1) ^24,25^ to dynamically adjust the spring force constants employed in the ANM, based on deep learning predictions of the physical contact probability between protein residues and membranes. Our MCP-ANM models provide a more accurate representation of the complex system of membrane proteins embedded in an anisotropic membrane environment, thereby avoiding reliance solely on simplistic boundary conditions. Consequently, these models are better suited for capturing the influence of the membrane on protein dynamics.

**Figure 1:**
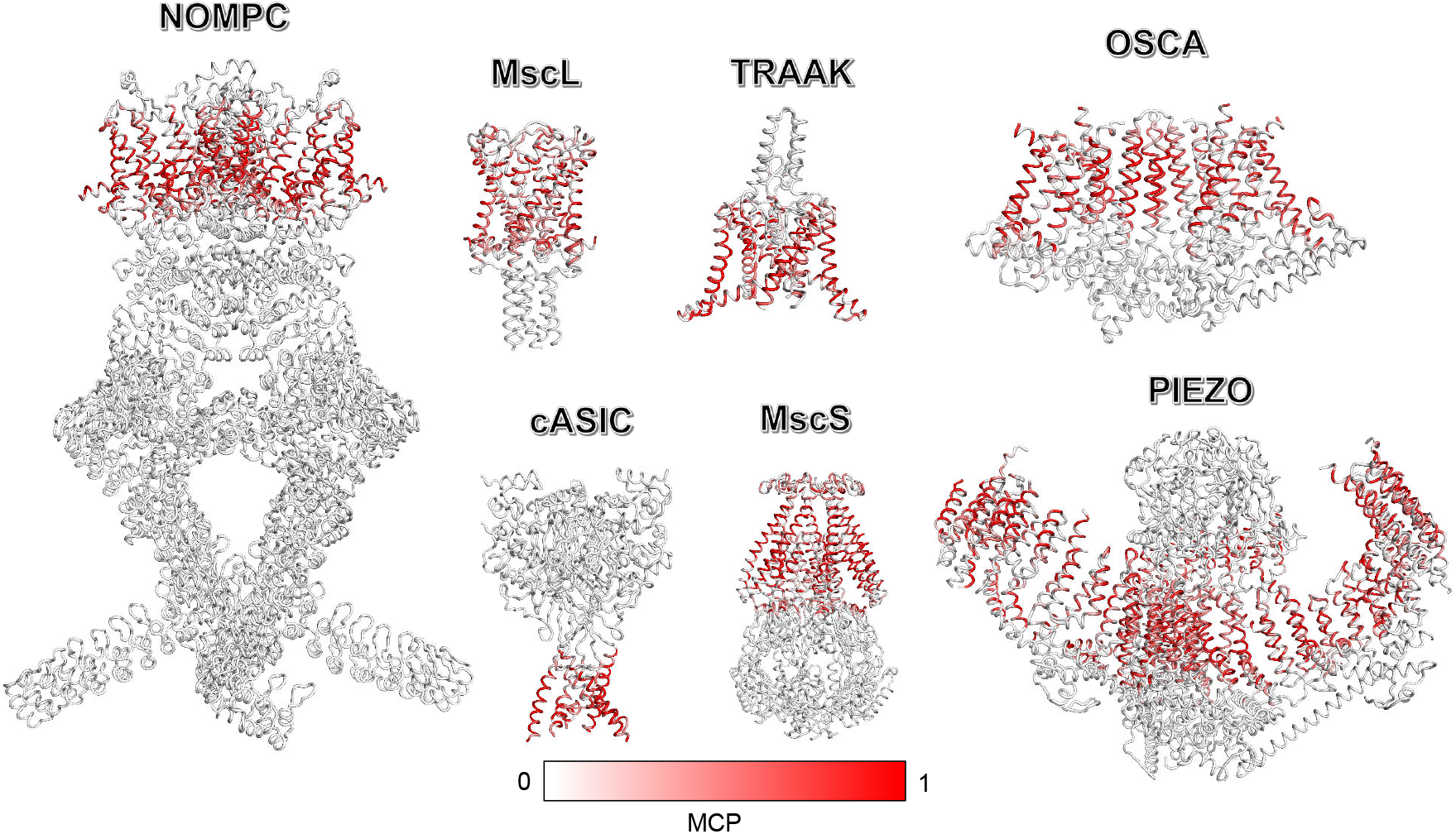
Structural representations of Mechanosensitive (MS) ion channels colored according to the predicted MCP. The protein structures are depicted as white tube models, with predicted MCP highlighted in red. The intensity of the red color indicates the likelihood of the region being in contact with the membrane, with more intense red signifying higher contact probability.

To further explore the gating mechanisms of MS ion channels, we combined MCP-based ANM with perturbation response scanning (PRS), a method that applies systematic perturbations to residues and evaluates the resulting conformational responses. ^26^ By extending PRS to incorporate MCP, we applied targeted mechanical forces to membrane-contacting or tethered residues of MS ion channels. This approach accurately simulates various gating mechanisms, including those induced by ‘force from lipids’ and ‘force from tether’. This integration enables a comprehensive analysis of the structural and functional dynamics of MS proteins within diverse mechanical environments. Our study highlights the efficacy of MCP-based ANM models in capturing the unique dynamics of MS ion channels and its potential to advance the understanding of mechanosensation.

## Results

### Framework Overview: MCP-ANM and MCP-mANM Models, and Integration with PRS

To incorporate the effects of the membrane environment into protein dynamics, we propose two modified Anisotropic Network Models (ANMs): MCP-ANM and MCP-mANM. In the MCP-ANM, nodes are defined as the C_*α*_ atoms of residues, and edges are established between node pairs within a defined cutoff distance. In the MCP-mANM, each pair of residues is connected by an edge, utilizing an exponential decay kernel function to represent residue interactions. ^27^ In both models, the uniform spring constant *γ* used in the vanilla ANM is replaced by direction-dependent spring constants based on the MCP for each residue. Residues with MCP values exceeding a defined threshold, which are considered membrane-contacting residues, are assigned modified spring constants to reflect the resistance exerted by the membrane during protein conformational changes (see ‘Methods’ section for details).

MCP is a predictive metric that quantifies the likelihood of direct interactions between amino acid residues and the acyl chains of lipid molecules. ^24,25^ It provides a complementary perspective to solvent accessibility, particularly for membrane proteins, whose surface hydrophobicity plays a crucial role. MCP values can be efficiently predicted from amino acid sequences using machine learning models trained on the MemProtMD datasets. ^28,29^ Incorporating predicted MCP values into ANM not only introduces membrane-specific constraints but also captures the intrinsic mechanical anisotropy of the membrane environment. Consequently, MCP-based ANM models may provide a more realistic representation of the constraints imposed by the membrane, enabling accurate simulations of protein dynamics in lipid bilayers. This approach also allows for detailed investigations of membrane protein behaviors in diverse mechanical environments, as demonstrated in subsequent sections.

To investigate the gating mechanism of mechanosensitive (MS) ion channels, we applied the MCP-based ANM framework in combination with the Perturbation Response Scanning (PRS) method^26^ to simulate mechanical stimulation. This integrated approach enabled a systematic analysis of how mechanical stress influences the structural dynamics and activation processes of MS channels. Notably, when coupled with PRS, the MCP-mANM framework tends to slightly relax the densely packed intra-protein connections. This subtle adjustment facilitates pore expansion under mechanical stress. In contrast, MCP-ANM, although effective in predicting flexibility (as will be demonstrated in the next section), maintains overly rigid internal connections, which can limit the conformational changes of pore in response to PRS-applied forces. As our computational results indicated that MCP-mANM more effectively captured the mechanical gating behavior of MS channels, it was selected for further investigation. The overall workflow of this study is illustrated in Figure 2.

**Figure 2:**
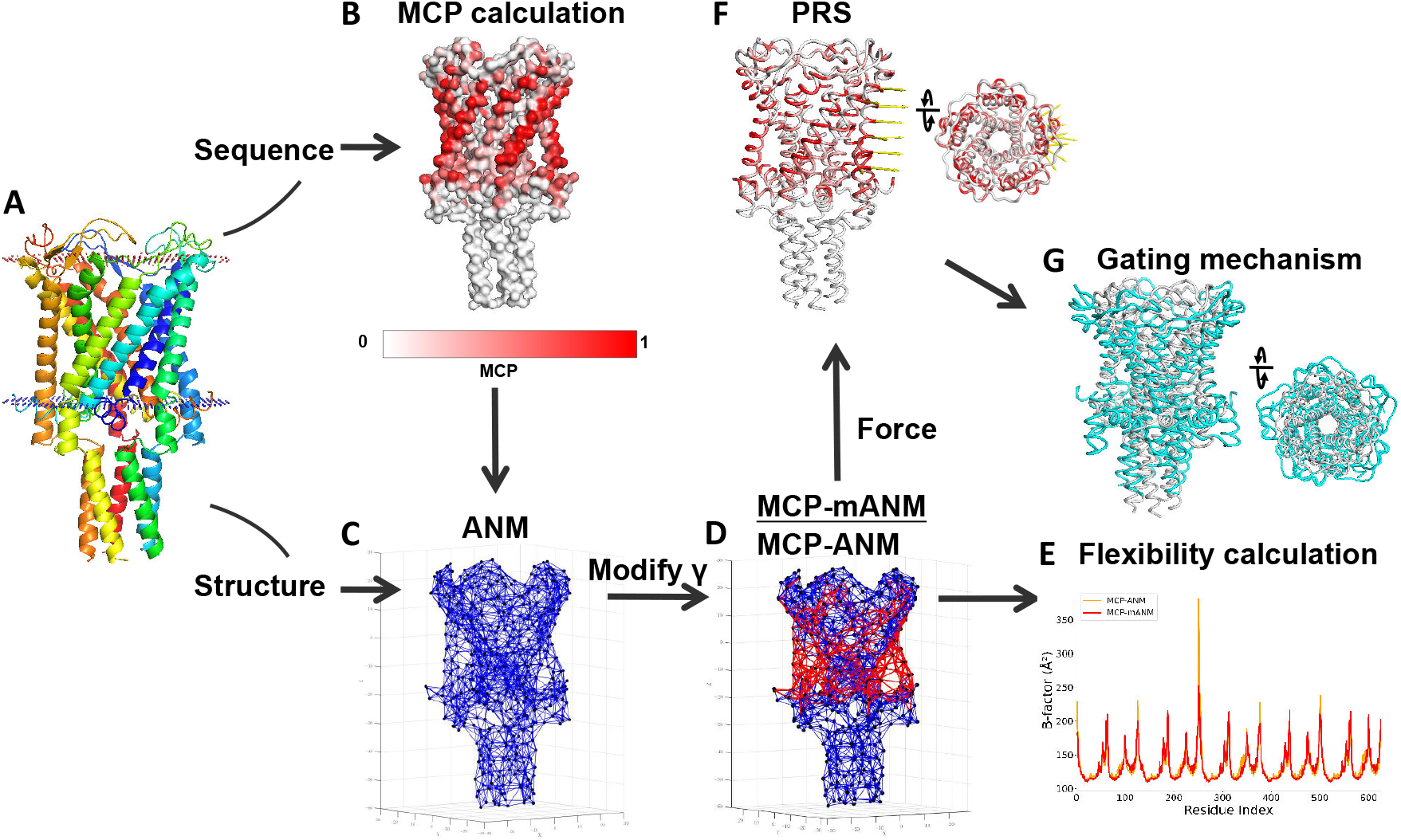
Workflow of combining MCP-based ANM and PRS to investigate the gating mechanisms of mechanosensitive ion channels. (A) Experimental structure of a MS protein (MscL as an example). (B) Predicted MCP values represented on the protein structure surface. (C) The vinilla ANM model with uniform spring constants (blue network). (D) Transition to the MCP-based ANM model, where spring constants are adjusted based on MCP values. (E) Flexibility profiles calculated using both MCP-ANM and MCP-mANM approaches. (F) PRS analysis using the MCP-mANM model with forces (yellow arrows) simulating membrane tension. For clarity, only the forces on one subunit are shown. (G) The resulting structural transition from closed (gray) to open (cyan) state.

Despite their structural and functional diversity, MS ion channels universally exhibit a gating mechanism that is activated by mechanical stimuli, leading to the opening of the pore and allowing for ion flux. Two primary models have been proposed to explain the gating mechanisms of MS ion channels:

1. Membrane Tension Model (‘force-from-lipids’ model): This model posits that mechanical tension or stress imbalance within the lipid bilayer itself triggers conformational changes in the embedded MS channel. When the bilayer is stretched, forces are transmitted directly to the channel through lipid-channel interactions, inducing structural re-arrangements that open the pore.
2. Spring-like Tether Model (‘force-from-tether’ model): In this model, MS ion channels are directly linked to intracellular or extracellular structural elements, such as the cytoskeleton or extracellular matrix, via spring-like tethers. Mechanical forces acting on these tethers induce conformational changes that propagate through the channel, leading to opening of the pores.

Combining MCP-mANM with PRS, we conducted simulations to examine how these two distinct force-transmission mechanisms influence the gating behavior of various MS ion channels. Specifically, we applied forces that mimic lipid bilayer tension or exert pushing forces along the tether structures to analyze the structural dynamics and transitions to the open state. The results indicate that these two models contribute differently to the gating process, depending on the channel type, its structural characteristics, and its interactions with the surrounding environment, which will be discussed in detail below.

### Theoretical B-Factor Calculation

B-factors, also known as the Debye–Waller factors, quantify the local flexibility of molecules and are commonly utilized to assess the dynamic properties of proteins. ^30,31^ We used Pearson’s correlation coefficient (PCC) values between theoretical and experimental B-factors to evaluate the performance of the ANM, imANM, MCP-ANM and MCP-mANM models for different types of membrane proteins collected from the MemProtMD database. ^28^ The results, presented in Figures 3A, 3B, and 3C, demonstrate that all three methods, MCP-ANM, MCP-mANM, and imANM, consistently outperform vanilla ANM in predicting B-factors across different types of membrane proteins, including ABC transporters, aquaporins, G protein-coupled receptors (GPCRs), ligand-gated ion channels, mechanosensitive ion channels, and outer membrane proteins. This suggests that accounting for membrane effects is crucial for accurately modeling the flexibility of membrane proteins. As shown in Figure 3D, MCP-ANM and MCP-mANM exhibit highly comparable performance in B-factor prediction, with MCP-ANM performing slightly better overall. Furthermore, Figures 3E and 3F compare the performances of MCP-ANM and MCP-mANM with that of imANM, illustrating the superior performance of MCP-based ANMs over imANM. These findings underscore that MCP-based ANMs not only capture the flexibility of membrane proteins more effectively than vanilla ANM and imANM but also demonstrate robustness and generalizability across diverse membrane protein types. By implicitly incorporating hydrophobic interactions at the lipid interface and the structural constraints imposed by membrane embedding, MCP-based ANMs provide a more accurate and comprehensive approach to modeling membrane protein dynamics compared to the vanilla ANM and imANM models.

**Figure 3:**
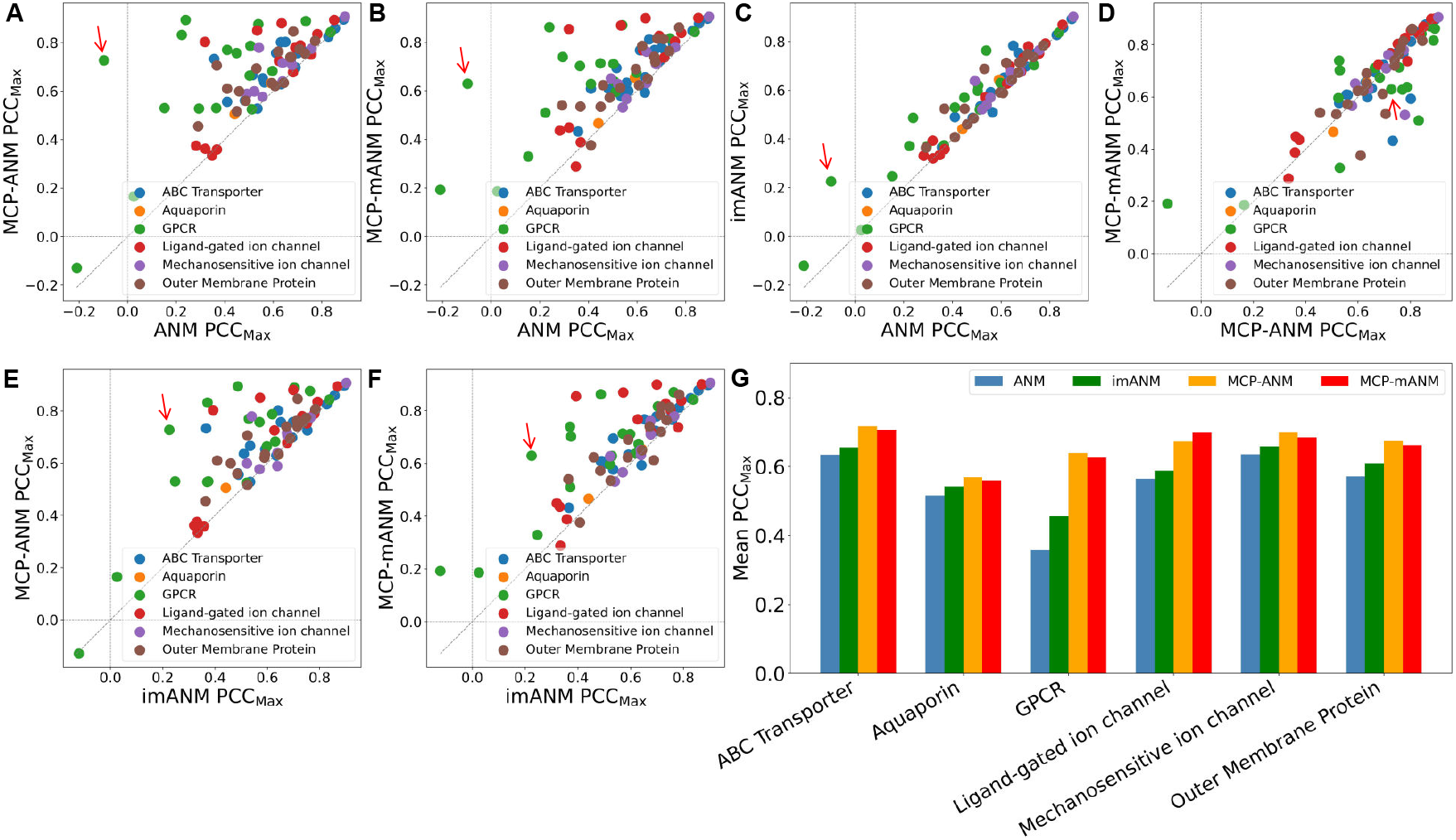
PCC values between experimental and theoretical B-factors calculated by four ANM models (ANM, imANM, MCP-ANM and MCP-mANM) for different types of membrane proteins. (A-F) Scatter plots showing the maximum PCC values for (A) ANM vs. MCP-ANM, (B) ANM vs. MCP-mANM, (C) ANM vs. imANM, (D) MCP-ANM vs. MCP-mANM, (E) imANM vs. MCP-ANM, (F) imANM vs. MCP-mANM. (G) Bar chart depicting the mean PCC values averaged by membrane protein type for ANM, imANM, MCP-ANM and MCP-mANM. The specific values of the mean PCCs are provided in Table S1.

As illustrated in Figure 3G, the most significant improvements are observed for GPCRs (G protein-coupled receptors), which are characterized by seven transmembrane helices and play a fundamental role in nearly all physiological processes. ^32,33^ Vanilla ANM models, which treat proteins as isolated systems, fail to account for the influence of the membrane, leading to discrepancies between theoretical and experimental B-factors. In some cases, the predictions even yield negative PCC values, indicating poor or even inverse correlations. For example, in the case of the Sphingosine 1-phosphate receptor 1 (the point in Figures 3A-3F, highlighted with a red arrow, with its structure shown in Figure 4A, PDB ID: 7VIE), the PCC value obtained with vanilla ANM was -0.098, emphasizing the model’s inadequacy in capturing membrane-protein interactions. imANM slightly improves the PCC to 0.255, but still falls short of capturing the true dynamics. In contrast, MCP-based ANMs improved PCC to 0.727 (MCP-ANM)/0.629 (MCP-mANM) by incorporating membrane-specific constraints into the modeling process, leading to a more realistic representation of the dynamics of GPCR. Figure 4B compares the predicted B-factors from the vanilla ANM, imANM, MCP-ANM and, MCP-mANM models with experimental data. The blue curve represents predictions from the vanilla ANM, which remain nearly flat and do not reflect experimental fluctuations (black curve). The green curve from imANM shows some improvement, but it still lacks accuracy. In contrast, the orange and red curves from MCP-ANM and MCP-mANM predictions align closely with the experimental values, particularly in capturing the peaks and troughs corresponding to the protein’s structural dynamics.

**Figure 4:**
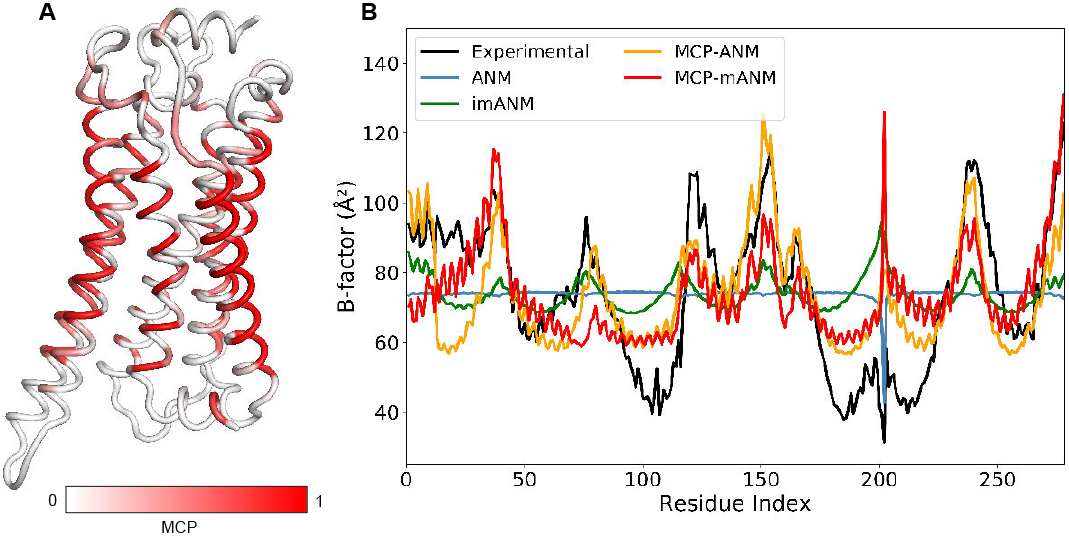
(A) Structure of Sphingosine 1-phosphate receptor 1 (PDB ID: 7VIE), colored according to the predicted MCP of residues. (B) The B-factor curves from experimental data and theoretical predictions calculated by four ANM models.

For the MS ion channels of particular interest in this study, MCP-based ANM also shows improvements in the prediction of B-factor, as shown in Table 1. The table summarizes the optimal PCC values for various MS ion channels modeled using four ANM models (the results including the optimal parameters and PCC for other membrane proteins are shown in Table S2). For example, the PCC of TRAAK (PDB ID: 4WFF) increases dramatically from 0.541 with vanilla ANM to 0.779 with MCP-ANM. Similarly, the PCC of PIEZO1 (PDB ID: 5Z10) improves from 0.522 to 0.627 with MCP-mANM. These enhancements underscore the capacity of MCP-based ANMs to more accurately model the membrane environment by accounting for critical interactions between proteins and the membrane, an aspect that is inadequately addressed by vanilla ANM models. Compared to imANM, MCP-based ANMs also show superior performance in most cases (68 out of 78 for MCP-ANM and 71 out of 78 for MCP-mANM, Tables 1 and S1). This suggests that MCP-based ANMs are more effective than imANM in capturing the influence of membranes on the dynamics of membrane proteins.

**Table 1:** Comparison of PCC_max_ Values from Different ANM Models for Mechanosensitive Ion Channels.

| Name | PDB ID | ANM | imANM | MCP-ANM | MCP-mANM |
| --- | --- | --- | --- | --- | --- |
| MscL | 2OAR | 0.556 | 0.570 | **0.577** | 0.566 |
| MscS | 6PWN | 0.639 | 0.639 | **0.640** | 0.632 |
| cASIC1 | 4NYK | 0.678 | 0.676 | 0.708 | **0.710** |
| OSCA1.2 | 6MGV | 0.896 | 0.902 | **0.907** | 0.905 |
| NOMPC | 5VQK | 0.757 | 0.765 | 0.775 | **0.779** |
| PIEZO1 | 5Z10 | 0.522 | 0.522 | 0.599 | **0.627** |
| TREK1 | 6CQ6 | 0.492 | 0.638 | 0.589 | **0.651** |
| TREK2 | 4XDJ | 0.639 | 0.675 | 0.716 | **0.761** |
| TRAAK | 4WFF | 0.541 | 0.540 | **0.779** | 0.531 |

$PCC_{\max}$ represents the highest Pearson correlation coefficient obtained for each method. The best-performing model for each protein is shown in bold.

By restricting unfavorable motions and incorporating membrane anisotropy, MCP-based ANMs enhance the accuracy of B-factor predictions and more effectively represent the dynamics of membrane proteins. However, the rigid internal connections present in MCP-ANM limit its applicability to large conformational changes induced by membrane tension. In contrast, the MCP-mANM offers a more flexible structure and a more accurate representation of large conformational changes, and was therefore chosen for subsequent analyses.

### Semiquantitative Analysis of known MS ion channels

To validate the accuracy of our predictions regarding the gating behaviors of MS channels, we applied normalized force vectors to a subset of MS channels whose gating mechanisms depend on pore dilation. We then quantitatively evaluated changes in the diameter of the gate, a critical indicator of gating behavior.

Figure 5 illustrates the progressive structural deformations of the MS channels in response to increasing membrane tension. Notably, under a certain membrane tension, PIEZO exhibits the most pronounced pore dilation, characterized by an outward displacement of the V2476 region, which aligns with its high mechanosensitivity. MscS exhibits moderate pore dilation, with an evident displacement of residue A102, while MscL shows relatively minor pore expansion, indicated by a slight displacement of residue L17. These findings highlight the varying mechanical sensitivities among the MS channels examined.

**Figure 5:**
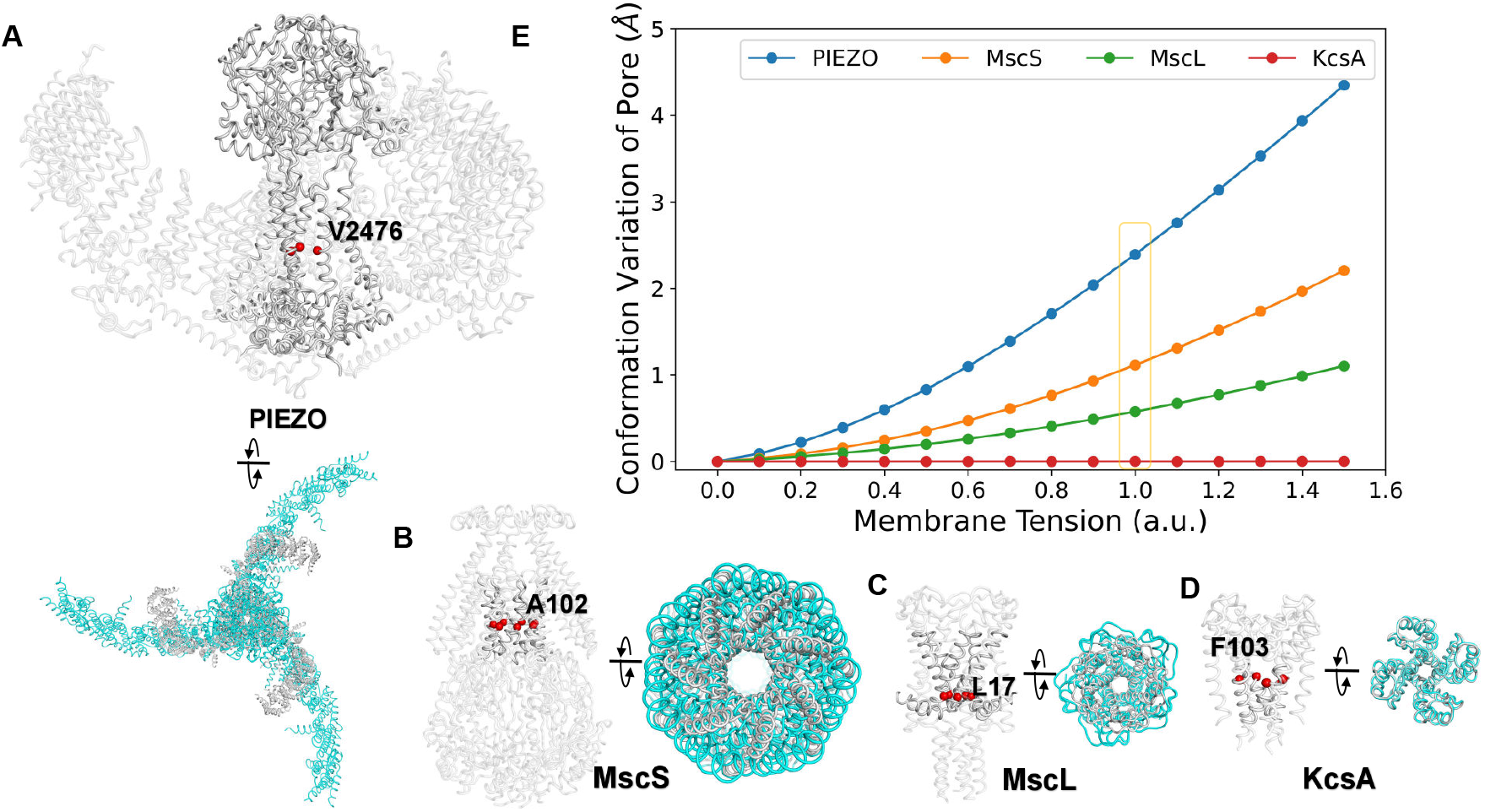
Response of MS channels to applied forces. In panels A–D, the gray structure represents the initial state, while the cyan structure shows the conformation after applied membrane tension (corresponding to the yellow-boxed point in panel E). Structural model of (A) PIEZO, (B) MScS, (C) MSCL, and (D) KcsA, with the pore gate residues used for analysis highlighted in red: V2476 (PIEZO), A102 (MScS), L17 (MSCL), and F103 (KcsA). (E) Comparison of pore conformation variation (diameter change at the gate) in the four ion channels under varying membrane tension (a.u.), showing PIEZO (blue) with the largest conformational change under certain membrane tension, followed by MScS (orange), MSCL (green), and KcsA (red).

For a consistent comparative analysis of the dilated structures of these channels, we selected a representative deformed structure for each channel at a tension level of 1.0 a.u. where significant conformational changes occur, as shown in Figures 5A-5C. At this tension level, each channel exhibits clear structural alterations, reinforcing the link between the applied membrane tension and gating behavior. Moreover, the calculated changes in the diameter of the pores at the gate (ΔR) increase with the applied membrane tension, and the relative sensitivity, derived from the slopes of the corresponding curves, follows the order PIEZO > MscS > MscL, consistent with experimental observations. ^34^ This agreement highlights the efficacy of our MCP-mANM model combined with the PRS method in capturing the mechanical sensitivity and gating behaviors of MS channels.

To validate the specificity of our model, we used the bacterial potassium channel KcsA (K^+^ channel of Streptomyces lividans, PDB ID: 1BL8) ^35^ as a negative control, as it is known to be insensitive to membrane tension. As illustrated in Figure 5, KcsA exhibited negligible conformational variation across all levels of applied membrane tension, demonstrated by the minimal changes in distance at its narrowest constriction site near residue F103. ^36^ The absence of a response in KcsA, in contrast to the MS channels, supports that the combination of MCP-mANM with the PRS method is capable of specifically detecting force-induced gating behaviors in MS ion channels.

However, it should be noted that due to the coarse-grained nature of the model, atomic-level details, such as precise pore sizes, remain beyond its scope. Consequently, the analysis is semiquantitative and primarily focuses on the overall trends at conformational changes rather than absolute values.

### Membrane Tension-Gated Ion Channels: Case Studies on MscS and PIEZO

To observe the gating transitions of MS proteins under mechanical stimuli and further demonstrate the applicability of the MCP-mANM model within the membrane tension-gated model, we selected MscS ^37^ and PIEZO ^38^ channels as representative examples. Both of these MS ion channels exhibit distinct gating behaviors when subjected to membrane tension. For further analysis, we selected conformations that exhibit an approximate 1 Å increase in pore radius: that is, the PIEZO structure under a membrane tension of 0.6 a.u. and the MscS structure at 0.9 a.u., as shown in Figure 5. This selection ensures meaningful conformational changes while minimizing the risk of excessive structural distortion caused by overstretching.

MscS undergoes significant conformational rearrangements during gating in the presence of membrane tension. ^39^ Experimental studies, including cryo-electron microscopy (cryo-EM), have identified two primary structural transitions that characterize MscS gating in response to mechanical stress: ^40^

1. Pore opening, which represents the conformational transition of the channel from a closed state to an open state, facilitating ion flow. This transition is highly dynamic and allows MscS to function as a sensor and regulator in cellular mechanotransduction pathways.
2. Rearrangements of transmembrane helices, which are essential for accommodating mechanical strain induced by membrane tension. Specifically, structural studies and computational simulations showed that, upon the application of tension, the transmembrane helices of MscS rotate outward, causing pore dilation.^40,41^ This expansion allows for a greater flow of ions through the pore.

Our simulations successfully captured the transmembrane helical rearrangements of MscS under tension. As shown in Figure 6A, the MscS structure is colored according to the membrane contact probability (MCP), highlighting regions with direct membrane interactions. Figure 6B illustrates the forces applied on one of the MscS chains, indicated by yellow arrows. Figure 6C provides a structural comparison of MscS before (gray) and after (cyan) the application of membrane tension, revealing significant conformational changes. Figures 6D and 6E further detail these changes: Figure 6D shows the rotation of the transmembrane helices in response to applied tension, while Figure 6E offers a top view of the structural changes, highlighting the helical rearrangements and the dilation of the pore. Figure 6F presents a zoomed-in view of the pore-lining helices, emphasizing their repositioning under force, and Figure 6G illustrates a top view of the pore region, showing the radial displacement and further dilation of the channel. The simulation results clearly demonstrate the rotation of the transmembrane helices, which closely aligns with experimental cryo-EM data. ^40^ This finding validates the reliability and accuracy of our model in reproducing the gating dynamics of MS ion channels at the molecular level.

**Figure 6:**
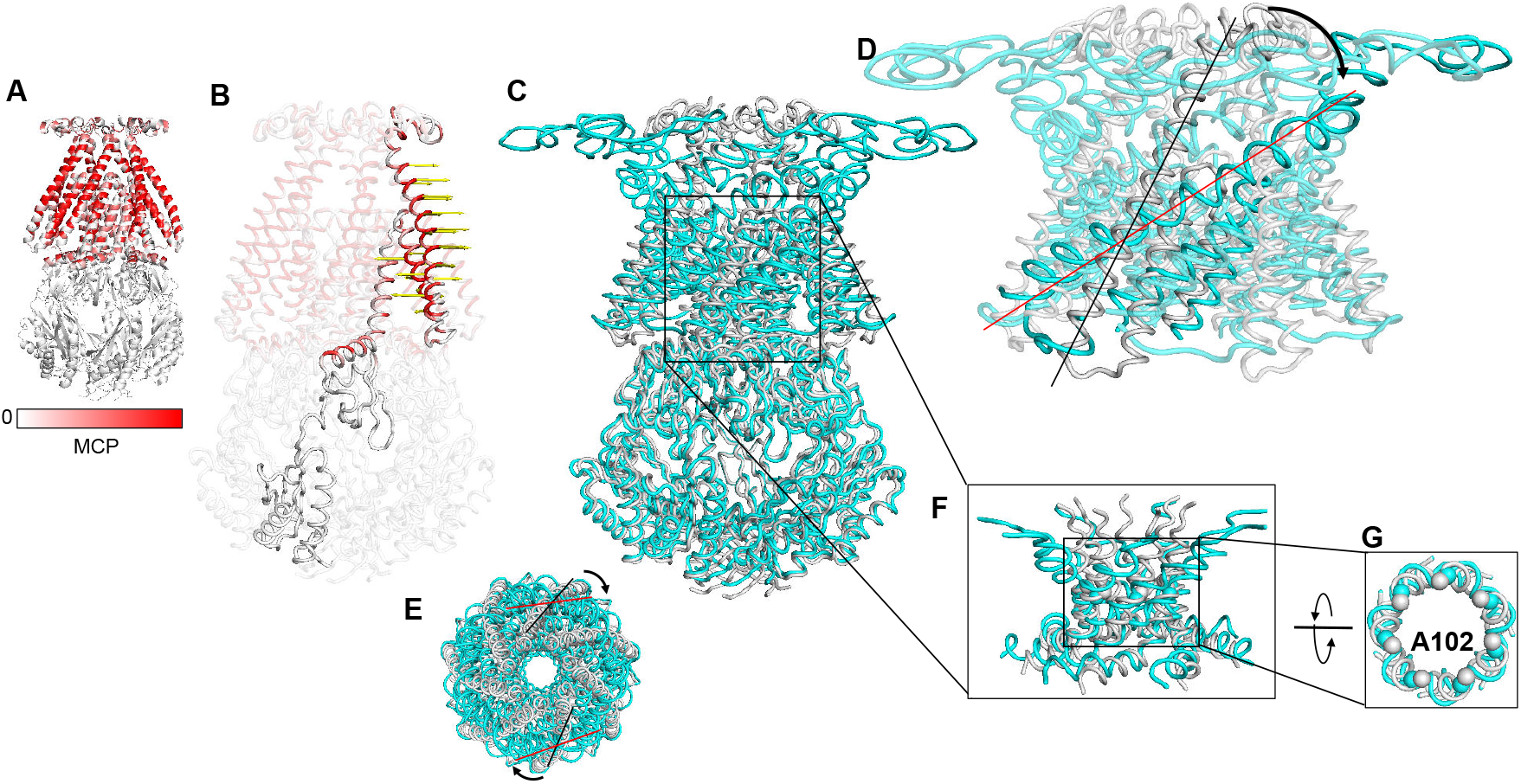
Structural Changes in MscS Under Membrane Tension. (A) MscS structure colored according to MCP. (B) The applied force on MscS is represented by a yellow arrow on one of the chains. (C) Structural comparison of MscS before (gray) and after (cyan) the application of membrane tension in our calculation. (D) Rotation of transmembrane helices in response to applied tension. (E) Top view showing the rotation and structural changes of MscS upon membrane tension, with the N-terminal of 20 residues omitted for clarity of the helical movements.. (F) Zoom-in on the pore-lining helices, showing their repositioning under force. (G) Top view of the pore region, illustrating the radial displacement and dilation of the channel. The spheres represent residue A102.

Another representative case is PIEZO, an MS ion channel that plays a pivotal role in the cellular response to mechanical stimuli such as membrane tension. PIEZO channels are involved in various physiological processes, including touch sensation, blood pressure regulation, and organ development.^42^ These channels are characterized by their unique structural design, which includes a large curved architecture that allows significant flexibility in response to mechanical forces. ^43^ Upon membrane tension, PIEZO undergoes large-scale conformational changes, facilitating its gating process and enabling ion permeation.^38^ Unlike MscS, the gating mechanism of PIEZO is driven by its distinctive curvature and the flexibility of its structure. Upon the application of membrane tension, PIEZO’s highly curved, dome-like architecture flattens, transitioning to a more planar form.^38^ This structural reorganization is accompanied by substantial rearrangements in the protein domains. Specifically, the cap-like domains at the apex of the protein rotate outward, leading to the widening of the central pore. ^44^ This shape change is believed to facilitate the passage of ions, as the widened pore allows for greater ion permeability, consistent with experimental findings from cryo-EM studies. ^45^ Previous studies have provided snapshots of the PIEZO’s gating process, revealing that membrane tension directly induces the flattening of the PIEZO structure, facilitating its transition from a closed state to an open state. ^38,46^

In our simulations, we successfully reproduced the large conformational changes observed in the gating of PIEZO. Figures 7A and 7B show the residues in contact with the membrane and the force applied to one of the PIEZO chains, respectively. Under simulated membrane tension, the PIEZO channel undergoes a gradual flattening process, with the top “cap” regions rotating outward, as indicated by the red arrows in Figures 7C and 7D. This flattening process is essential for the gating mechanism. As shown in Figure 7C, the side view reveals the structural changes from a curved to a flattened state, as well as the outward of the cap regions, leading to the widening of the central pore. Figure 7D provides a top view that further demonstrates the complete domain rearrangements, particularly emphasizing the rotational changes. In addition, Figure 7E presents a zoomed-in view of the pore region, illustrating the conformational shift of pore-lining helices under tension, while Figure 7F offers a top view that depicts the radial displacement and dilation of the channel. These structural changes indicate the opening of the pore in response to membrane tension, aligning well with experimental observations and cryo-EM data. ^45^ These findings highlight the accuracy and predictive capability of our computational approach in capturing the large-scale conformational transformations that are integral to the gating mechanism of PIEZO.

**Figure 7:**
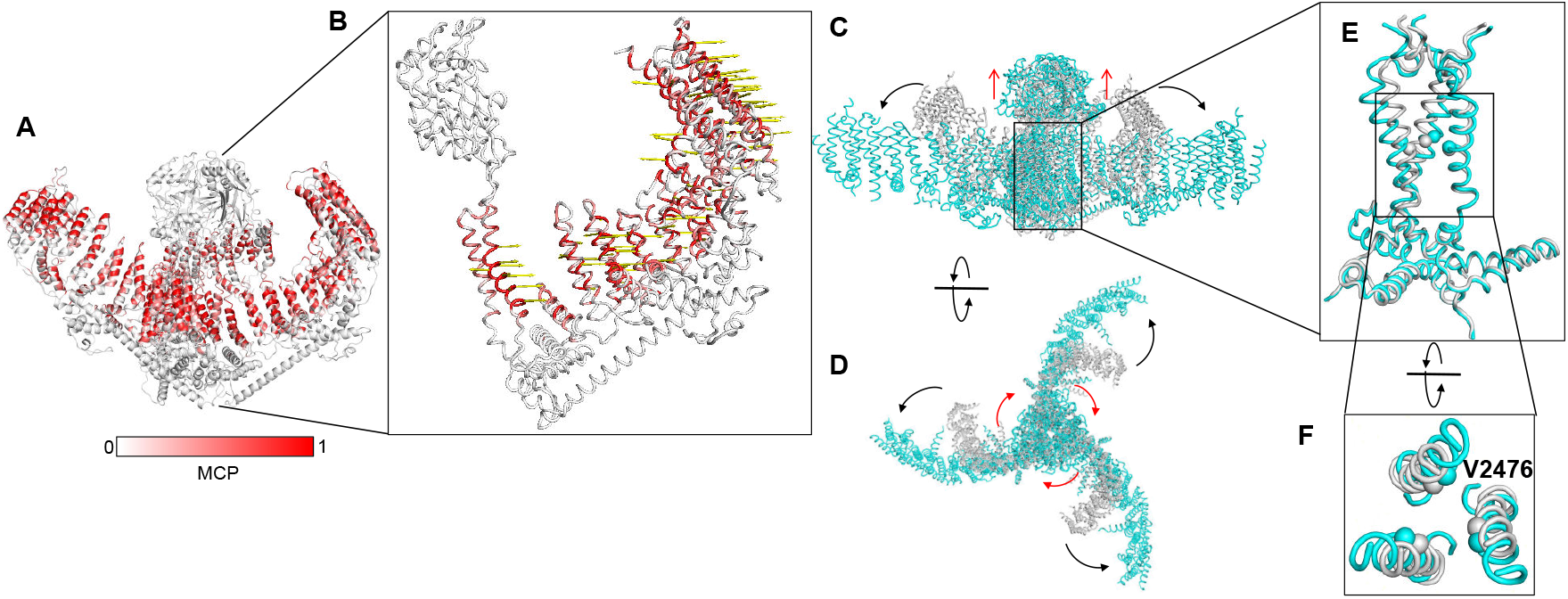
Structural changes of PIEZO1 under membrane tension in our calculation. (A) PIEZO1 structure colored according to the predicted MCP. (B) The forces applied on PIEZO1, depicted by yellow arrows on one chain . (C) Side view of the structural changes from a curved to a more flattened state and (D) corresponding top view. The red arrow indicates the changes of the top ‘cap’. (E) Zoom-in on the pore region, showing the conformational shift under tension. (F) Top view of the pore region, illustrating the radial displacement and dilation of the channel.

### Tether-Gated Ion Channels: NOMPC as an Example

To evaluate the applicability of our method to the tethered mechanosensation model, we selected NOMPC, a member of the transient receptor potential (TRP) ion channel family, ^47^ as a representative example. NOMPC is characterized by extended ankyrin repeat (AR) domains that serve as force sensors, linking external mechanical stimuli to the channel’s pore region. This structural configuration facilitates a “push-to-open” or “twist-to-open” gating mechanism, wherein mechanical forces applied to the AR domains are transmitted through the TRP domain to the pore-lining helices (S6), thereby inducing channel opening. ^15,16,48^ Previous studies have shown that compressing the AR domains results in clockwise rotation of the TRP domain, which leads to pore opening.^15,16,48^ Based on this, we applied upward pushing forces to the AR1 domain of NOMPC and analyzed the resultant conformational changes.

As illustrated in Figure 8, the application of an upward push force to the ankyrin repeat (AR1) domain of NOMPC resulted in a series of structural transformations that align with the tether-gating model. The structure shown in Figures 8A-8D correspond to the conformation obtained at a force magnitude of 0.03 a.u., as indicated along the x-axis in Figures 8E-8H. The initial mechanical stimulus induced compression within the ankyrin repeat (AR) domains (Figures 8A, 8D and 8E), resulting in a cascade of deformations that propagated to the TRP domain. This force transmission triggered a clockwise rotation of the TRP domain about the channel’s central axis (Figures 8C and 8G), which serves as a crucial intermediate step in converting linear mechanical force into rotational movement. The rotational motion of the TRP domain induces a progressive tilt in the S6 helices (Figures 8B and 8F), which comprise the channel pore, resulting in an increase in pore diameter (the C*α*-C*α* distance of opposite I1554 residues that form the narrowest part of the gate) (Figure 8H). This sequential propagation of force and deformation, originating from the AR domains to the TRP domain and ultimately to the pore-lining helices, underscores the intricate compress-twist coupling mechanism that drives mechanosensitive gating in NOMPC.

**Figure 8:**
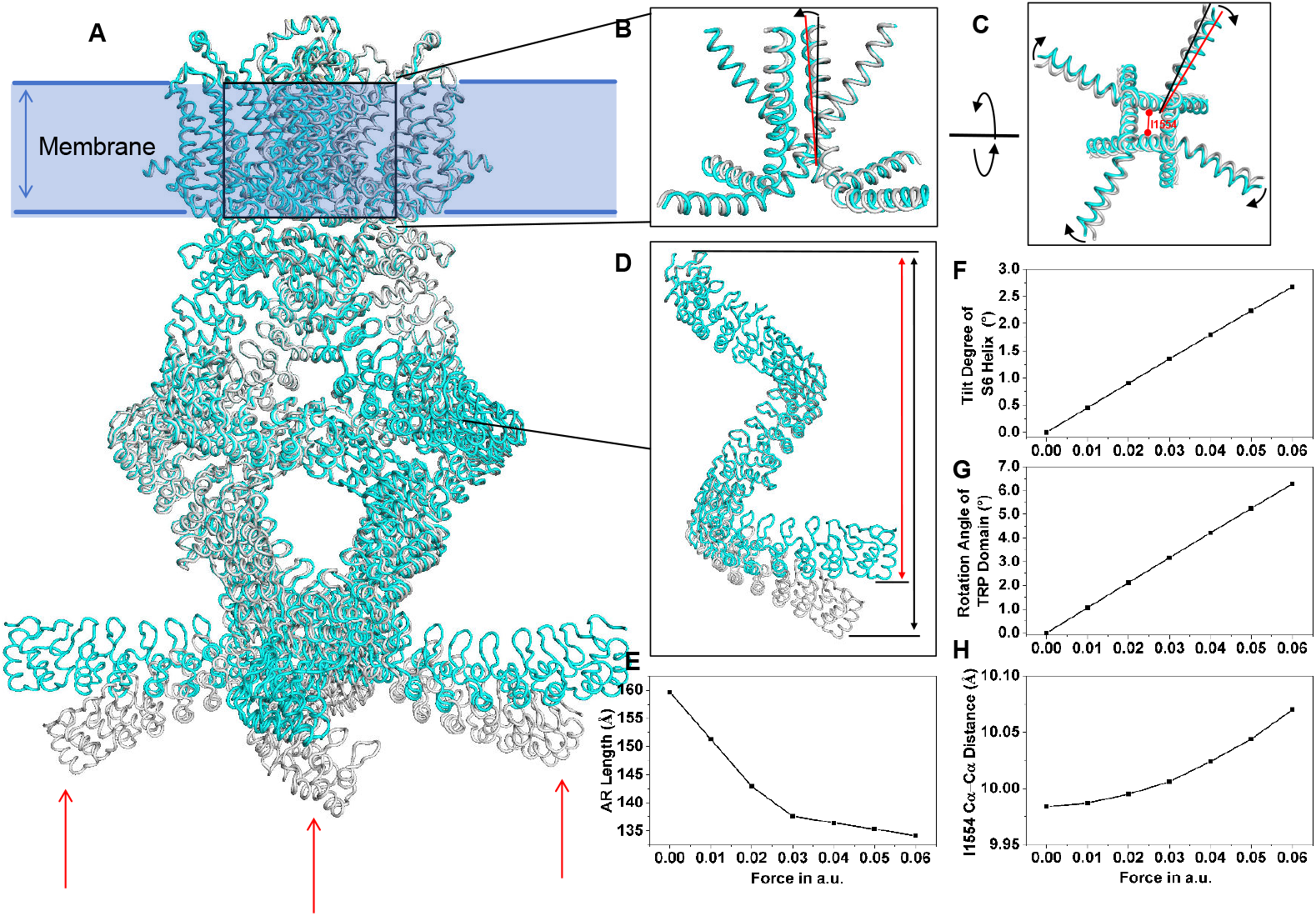
Structural changes during the push-to-open gating process of NOMPC. (A) Overall structural changes with the initial state shown in gray and the deformed state in cyan. The cell membrane is represented by a light blue strip. The red arrows indicate the direction of the applied forces. (B) The S6 helix slightly tilts downward. (C) The TRP domain rotates clockwise (viewed from the intracellular side) under the torque generated by the pushing force. (D) The length change of the AR spring under compression, with the initial state in gray and the compressed state in cyan. (E)-(H) The structural changes as a function of applied forces, illustrated by the length of the AR domain (E), the tilt angle of the TRP domain (F), the rotation angle of the TRP domain viewed from the intracellular side (G), and the pore size represented by the distance between two C_α_ atoms of the I1554 residues on the two opposite subunits (H).

These results underscore the critical role of AR domains in the gating mechanism of NOMPC. By functioning as initial force sensors, the AR domains transmit mechanical forces to the TRP domain, thereby facilitating pore opening. This supports the hypothesis that the coupling between AR and TRP domains is essential for NOMPC’s mechanosensitivity, especially regarding its structural adaptations in response to mechanical stimuli.^47^ Moreover, these results demonstrate the broader applicability of our approach, with both membrane tension- and tether-gating mechanisms being nicely accounted for.

## Discussion

The computational study of the large-scale dynamics of membrane proteins remains challenging, particularly when external stimuli such as mechanical forces are involved. Conventional coarse-grained methods often omit the influence of the membrane environment on proteins, making it difficult to accurately capture the dynamic properties of membrane proteins within their membrane surroundings. By integrating predicted MCP values into the Anisotropic Network Model (ANM), our study significantly enhances its predictive accuracy, establishing MCP-based ANM as a robust tool for investigating the dynamics of membrane proteins. This integration provides a more realistic representation of the behavior of membrane proteins under the influence of the membrane environment, enabling efficient analysis of their conformational flexibility.

The Perturbation Response Scanning (PRS) method based on MCP-mANM introduces a new approach to quantifying the mechanical force responses of mechanosensitive ion channels. In contrast to previous applications of the PRS method, which perturbed all residue positions to identify those critical for overall protein function and conformation, our strategy begins with known functionally significant residues located on the proteins. By applying perturbations or mechanical forces to these membrane-contacting or force-acted residues, we can efficiently simulate how these mechanical stimuli drive conformational changes. This approach shifts the focus from merely identifying key residues to exploring underlying functional mechanisms, thereby providing new insights into gating mechanisms and aligning closely with experimental observations.

Our method’s capacity to replicate actual membrane tension by applying forces to membrane-contacting residues enables it to capture the intrinsic dynamics of MS ion channels within their native environment. This capability is essential for accurately reproducing the gating mechanisms observed in experimental studies. For instance, our simulations of MscS and PIEZO channels successfully replicated their large-scale conformational changes in response to membrane tension, demonstrating the method’s accuracy and predictive power. Meanwhile, for tether-gated MS ion channels such as NOMPC, our simulations also revealed the gating processes and detailed conformational changes, further validating the robustness of our approach.

This versatility underscores the potential of our methodology to investigate the gating mechanisms of other unknown or less-characterized MS channels, offering valuable insights into their functional regulation and role in cellular physiology. In addition to these advantages, our method is computationally efficient and significantly faster than traditional MD simulations. All computational experiments were conducted on a workstation equipped with an Intel® Xeon® W-2255 CPU operating at 3.70 GHz (10 physical cores, 20 threads) and 64 GB of RAM. To ensure accurate and consistent timing, computations were executed in a single-threaded environment. We evaluated the scalability of our method using three protein structures of increasing size: MscL (PDB ID: 2oar, 625 residues), MscS (PDB ID: 6pwn, 1960 residues), and NOMPC (PDB ID: 5vkq, 5996 residues). The corresponding computation times were 3.89 seconds, 26.30 seconds, and 191.25 seconds for the MATLAB version, and 6.45 seconds, 77.89 seconds, and 790.14 seconds for the Python version, respectively. This speed enables the rapid elucidation of gating mechanisms for MS ion channels, establishing it as a powerful tool for high-throughput investigations.

Despite its strengths, our MCP-based ANM method has several limitations. First, as a coarse-grained approach, it lacks atomistic details, a common drawback of ANM-based models. This limitation may restrict its ability to accurately capture fine-scale molecular interactions, such as specific lipid-protein contacts. Second, the performance of the method depends on the accuracy of MCP calculations. Although MCP predictions are generally reliable, ^24,25^ errors in the MCP calculations may still impact the overall results, particularly for proteins that exhibit atypical membrane interactions. Nevertheless, MCP-based ANM remains a rapid and effective tool for elucidating the gating mechanisms of MS ion channels, providing valuable insights into membrane protein dynamics.

In summary, this study presents a new computational framework for investigating the dynamic properties of membrane proteins. The integration of MCP data into ANM and the refinement of PRS significantly enhance the predictive capabilities of these methodologies, thereby advancing the field of membrane protein dynamics studies. By providing a more realistic and efficient approach to studying MS ion channels, our methodology facilitates deeper insights into their functional mechanisms, supports high-throughput MS protein research, and offers potential applications in biomedical research.

## Methods

### Dataset

To evaluate the performance of MCP-based ANM for membrane proteins, we compiled a dataset of membrane proteins from the MemProtMD database, ^28^ a database generated through molecular dynamics simulations of membrane proteins with known structures. These proteins were classified according to the MemProtMD scheme, with an additional category for mechanosensitive (MS) proteins, resulting in a total of six distinct categories and 2,090 membrane proteins. The membrane protein structures were then selected based on the following criteria: resolution < 4.0 Å and exclusion of structures derived from NMR data. Ultimately, we obtained a total of 1,873 membrane proteins.

To minimize redundancy and enhance the diversity of the dataset, we conducted structure clustering using the Foldseek tool,^49,50^ ultimately selecting 78 representative structures. The clustering was carried out to ensure that the structural overlap among the chosen proteins did not exceed 90%, while the structural similarity (E-value) between the groups remained below A comprehensive list of the final selected structures is provided in Table S2.

### MCP prediction

The membrane contact probability (MCP) predictor was employed to quantify the likelihood of residue-membrane contacts with 6 Å as the distance cutoff, which requires only the protein sequence in FASTA format as input.^24,25^ For a given protein sequence, the MCP predictor outputs per-residue values on a [0,1] scale, with increasing values corresponding to stronger predicted contact probabilities with the lipid membrane. Recently, we developed a new version of the transformer-based MCP predictor, which uses language model embeddings as input features, offering significantly enhanced speed and accuracy. ^25^ In this study, we employ this updated version to perform the MCP predictions. The source code for our MCP predictor is accessible at https://github.com/ComputBiophys/ProtRAP-LM. An online computation server is available for http://www.songlab.cn/ProtRAP-LM/home/.

### ANM

The Anisotropic Network Model (ANM) is a simplified molecular dynamics approach utilized for the investigation of protein dynamics. In conventional ANM, a protein structure is represented as an elastic network composed of selected atoms (here C_*α*_). In this network, pairs of atoms/nodes that are within a specified cutoff distance (*r*_*c*_) are interconnected by springs, each characterized by a uniform force constant (. ^20^ The total potential energy of the network can be written as:

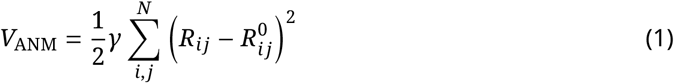

where *R*_*ij*_ and 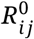 refer to the instantaneous and equilibrium distances between node *i* and *j* respectively. The dynamic properties of proteins are determined by a Hessian matrix ***H***, whose elements are 3 ×3 submatrices. The submatrix ***h***_*ij*_ is calculated as the matrix of second-order derivatives of the potential with respect to the Cartesian coordinates of the nodes. When *i* ≠ *j*, the corresponding ***h***_*ij*_ is:

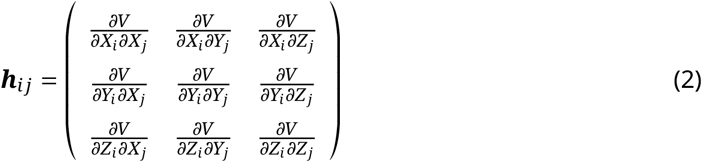

When *i* = *j*, the submatrix is: ***h***_*ij*_ = − ∑_*i≠j*_ ***h***_*ij*_

The dynamical properties can be determined using the pseudoinverse of the Hessian matrix, as detailed in the subsequent section regarding the calculation of B-factors.

### Multiscale ANM (mANM)

In the multiscale ANM (mANM) approach, interactions across multiple scales are typically modeled using different exponential decay kernel functions to accurately capture both short- and long-range couplings. ^27^ In our study, we adopted a simplified variant that employs only a single exponential decay kernel function. While this reduction in complexity limits the capability to model hierarchical interactions, it proves advantageous in our MCP-mANM (refer to the next section). This simplification not only streamlines parameter tuning but also better preserves the essential internal dynamics relevant to mechanosensitive gating. Specifically, it alleviates excessive rigidity within the protein interior that results from dense local packing, thereby enabling more realistic conformational transitions in response to mechanical stress.

In mANM, the interaction strength between nodes *i* and *j* is defined by:

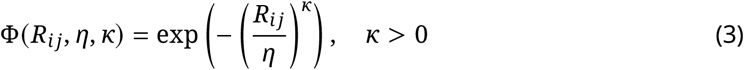

where *R*_*ij*_ is the distance between the *i* th and *j*th nodes, and the parameters *η* and *κ* control the decay extent.

When *i* ≠ *j*, the submatrix corresponding to the (single) kernel function is given by:

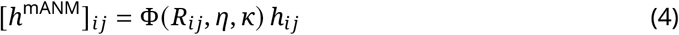

and for *i* ≠ *j*, the diagonal element is computed as:

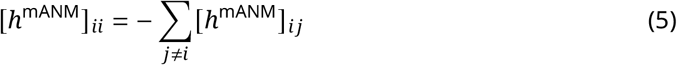

This formulation allows us to attenuate the contribution of strong near-range interactions directly through the kernel function, thereby reducing network stiffness and enabling the simulation of large-scale conformational transitions that are essential for mechanosensitive channel gating.

### MCP-ANM and MCP-mANM

We modified the vanilla ANM and mANM by replacing the uniform spring constant *γ* with direction-dependent spring constants *γ*_*x*_, *γ*_*y*_, *γ*_*z*_. For residues with MCP values exceeding a certain threshold, the new spring constants are adjusted to reflect the influence of the membrane environment, as shown in Eq. 6.

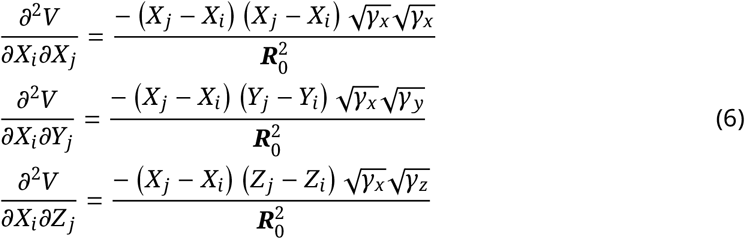

This modification enables the independent adjustment of force constants along each Cartesian coordinate, facilitating a more precise simulation of protein dynamics within the membrane environment. Specifically, the membrane’s resistance to protein motion is modeled through two scaling factors: *γ*_*x*_ = *γ*_*y*_ = s_1_ > 0: A scaling factor for radial motions (in the x- and y-directions), which captures the membrane’s resistance to lateral displacements of the protein, simulating the constraining effect of the lipid bilayer on the protein’s movement within the plane of the membrane. *γ*_*z*_ = s_2_ > 0: A scaling factor for motions normal to the membrane (along the z-direction), which accounts for the additional resistance experienced by the protein when attempting to move across the lipid bilayer, as the membrane is more resistant to vertical movement.

These modifications enable us to model the perturbations in protein dynamics that result from the physical properties of the membrane, offering a more precise representation of the behavior of membrane proteins in their natural environment.

### Calculation of B-factors based on ANMs

In ANMs, the dynamic properties can be expressed as:

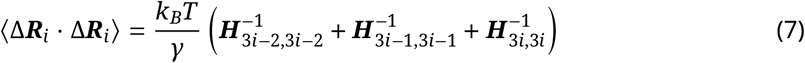

The pseudoinverse of the Hessian matrix can be decomposed as:

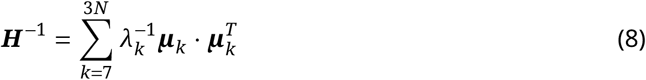

where *λ*_*k*_ and ***μ***_*k*_ are the *k* th eigenvalue and eigenvector of the Hessian matrix, respectively. According to Debye-Waller theory, the theoretical B-factor for the *i* th node can be calculated using the following expression:

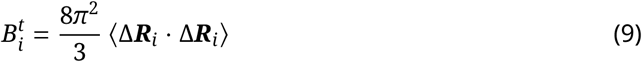

The Pearson correlation coefficient (PCC) is used to evaluate the degree of correlation between theoretical and experimental B factors, which is given by:

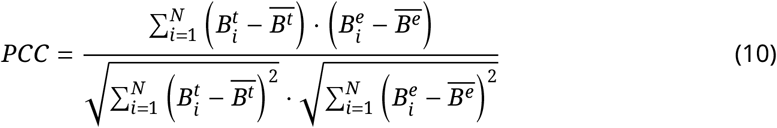

where 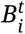 and 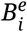 are the theoretical and experimental B-factors for the i-th node, respectively, and 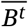 and 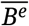 are their mean values. The PCC ranges from -1 to 1, with a value of 1 indicating perfect correlation and a value of -1 indicating perfect anticorrelation.

It should be noted that for the vanilla ANM, imANM, MCP-ANM, and MCP-mANM models, optimized parameters are determined to construct the corresponding ANM models by maximizing the PCC value between the theoretical and experimental B-factors for each protein. During this optimization process, the cutoff distance is systematically varied within the range of [8 Å, 20 Å] with a step size of 1 Å. In the case of imANM, the model defines the spring constants in the x-y plane as multiples of the spring constant in the z-direction, which is fixed at a value of 1 by default in the program. ^23^ Consequently, the multiplier for the x-y spring constant (relative to the z-direction) is varied across the values [1, 2, 4, 8, 16, 32, 64]. For the MCP-based ANM, the MCP cutoff parameter is systematically varied in the range of [0.1, 0.9] with a step size of 0.1, while the parameters s_1_ and s_2_ are varied in the range of [1, 2, 4, 8, 16, 32, 64]. For MCP-mANM, the parameter *η* varies across the range of [1, 8] and the parameter *κ* varies in the range of [1, 3] with a uniform step size of 1.

### Perturbation response scanning approach

The perturbation response scanning (PRS) approach^26^ based on linear response theory (LRT) ^51^ was developed to elucidate the allosteric properties of proteins. This method enables the calculation of the response of a residue *k* to a perturbation applied to another residue *i*. The 3*N*-dimensional vector Δ***R*** of node displacements in response to the exertion of a perturbation (3*N*-dimensional force vector ***F***) obeys Hooke’s law ***F*** = ***H***Δ***R***, where ***H*** is the 3*N* × 3*N* Hessian matrix in the Anisotropy Network Model (ANM) theory. The force exerted on the residue *i* is written as:

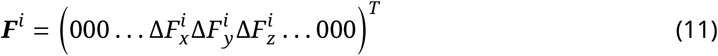

and the resulting response is:

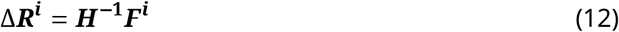

where Δ***R***^*i*^ is a 3*N*-dimension vector describing the displacements of all residues away from their equilibrium positions in response to the exerted force ***F***^*i*^, which are nonzero only for the three terms related to residue *i*.

PRS systematically applies forces to individual residues and records the linear response of the entire protein. This response is quantified by analyzing both the magnitude of residue displacements and their directionality. In this study, we extend the PRS method to investigate the responses of MS proteins to mechanical forces by applying perturbation force vectors to membrane-contacting or force-acted residues. This extension facilitates precise control over the direction and magnitude of the applied forces, enabling the simulation of mechanical impacts on MS proteins. For MS proteins with different gating mechanisms, such as “force-from-lipids” and “force-from-tether”, we employ tailored strategies to ensure a comprehensive analysis of all types of MS proteins.

For MS proteins operating under the “force-from-tether” mechanism, forces are applied directly to the tethered residues along a predefined axis (e.g. perpendicular to the membrane plane). For the “force-from-lipids” mechanism, forces are similarly applied to membrane-contacting residues. The applied force consists of two components:

1. Surface Normal Vector: The tension is aligned with the normal vector of the surface at the location of each residue.
2. Radial Vector Adjustment: To refine the direction of the force, the normal vectors are projected onto the X − Y plane. Subsequently, these vectors are further adjusted using radial vectors that are centered at the geometric center of the protein.

Residues with MCP > 0.9 were subjected to membrane tension. The code utilized in this study is publicly available on GitHub.

For MS proteins gated by membrane tension, we applied the same magnitude of force to each membrane-contacting residue of the ion channel examined.

## Data and code availability

The code will be available after peer review at https://github.com/ComputBiophys/MCP-ANM. For the purposes of review, the code is supplied as a supplementary file.

## Acknowledgements

This work was supported by the National Key R&D Program of China (2024YFA0916800), the Science Fund for Innovative Research Groups of the National Natural Science Foundation of China (T2321001), and the Postdoctoral Fellowship from the Peking-Tsinghua Center for Life Sciences. C.S. was supported in part by the Frontier Innovation Fund of Peking University Chengdu Academy for Advanced Interdisciplinary Biotechnologies.

## TOC Graphic

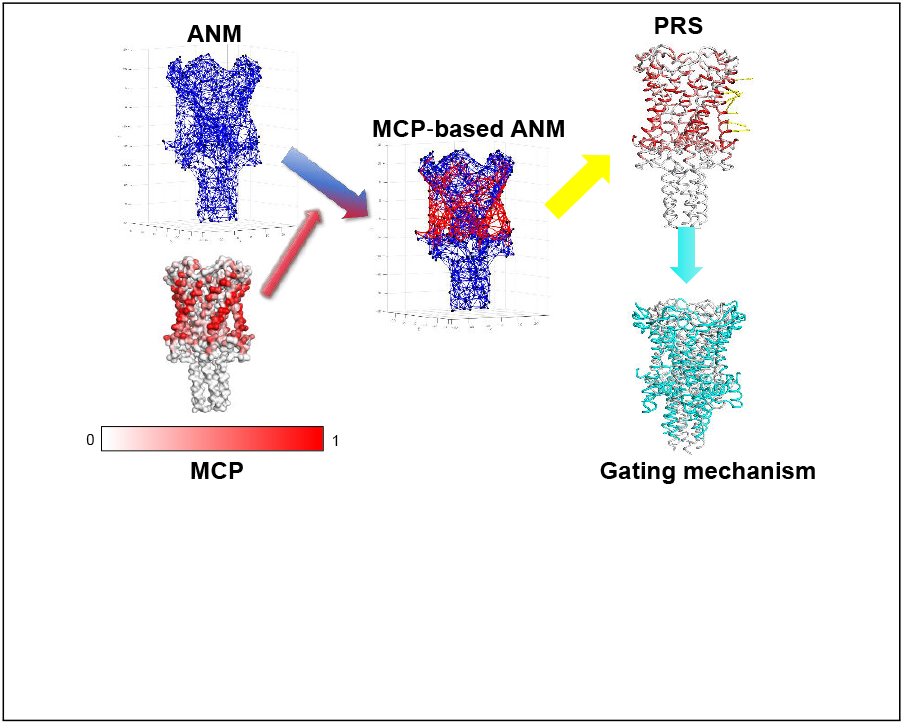

## Supporting Information

**Table S1:** Mean PCC_max_ values averaged by membrane protein types for ANM, imANM, MCP-ANM, and MCP-mANM.

| Membrane Protein Type | ANM | imANM | MCP-ANM | MCP-mANM |
| --- | --- | --- | --- | --- |
| ABC Transporter | 0.634 | 0.656 | **0.717** | 0.707 |
| Aquaporin | 0.516 | 0.542 | **0.569** | 0.560 |
| GPCR | 0.358 | 0.456 | **0.639** | 0.626 |
| Ligand-gated ion channel | 0.565 | 0.588 | 0.673 | **0.698** |
| Mechanosensitive ion channel | 0.636 | 0.659 | **0.699** | 0.685 |
| Outer Membrane Protein | 0.571 | 0.609 | **0.674** | 0.662 |

**Table S2:**
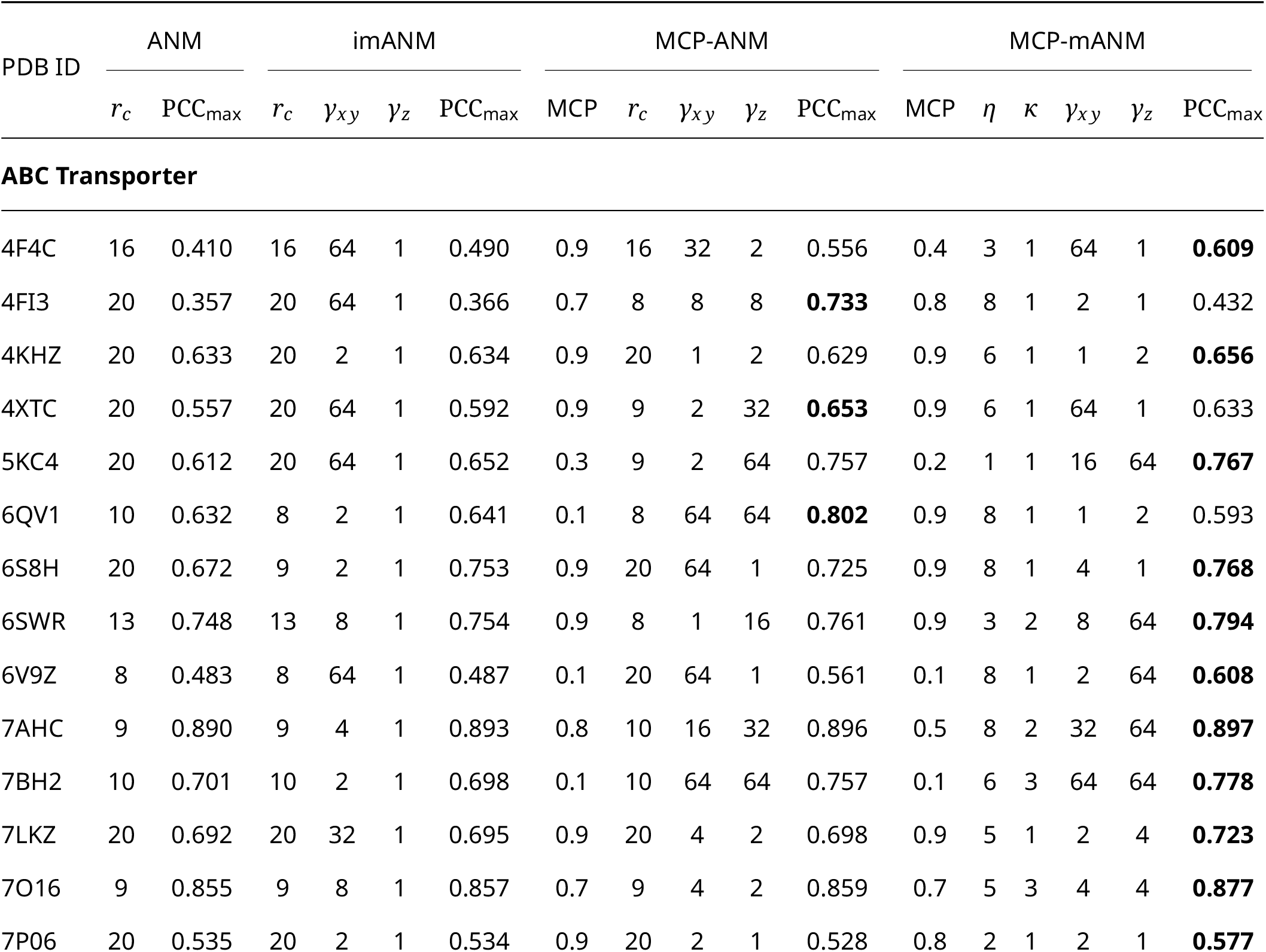
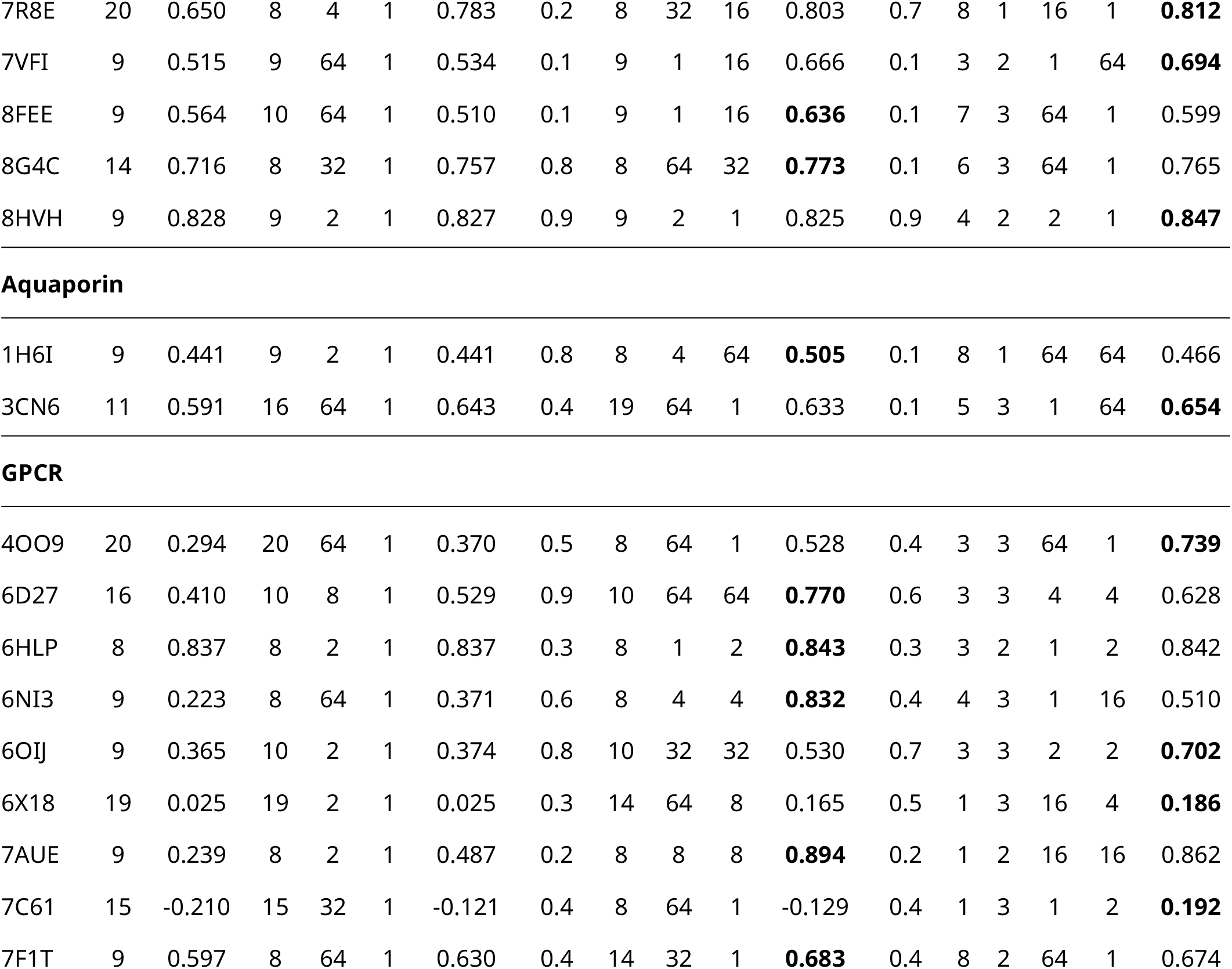
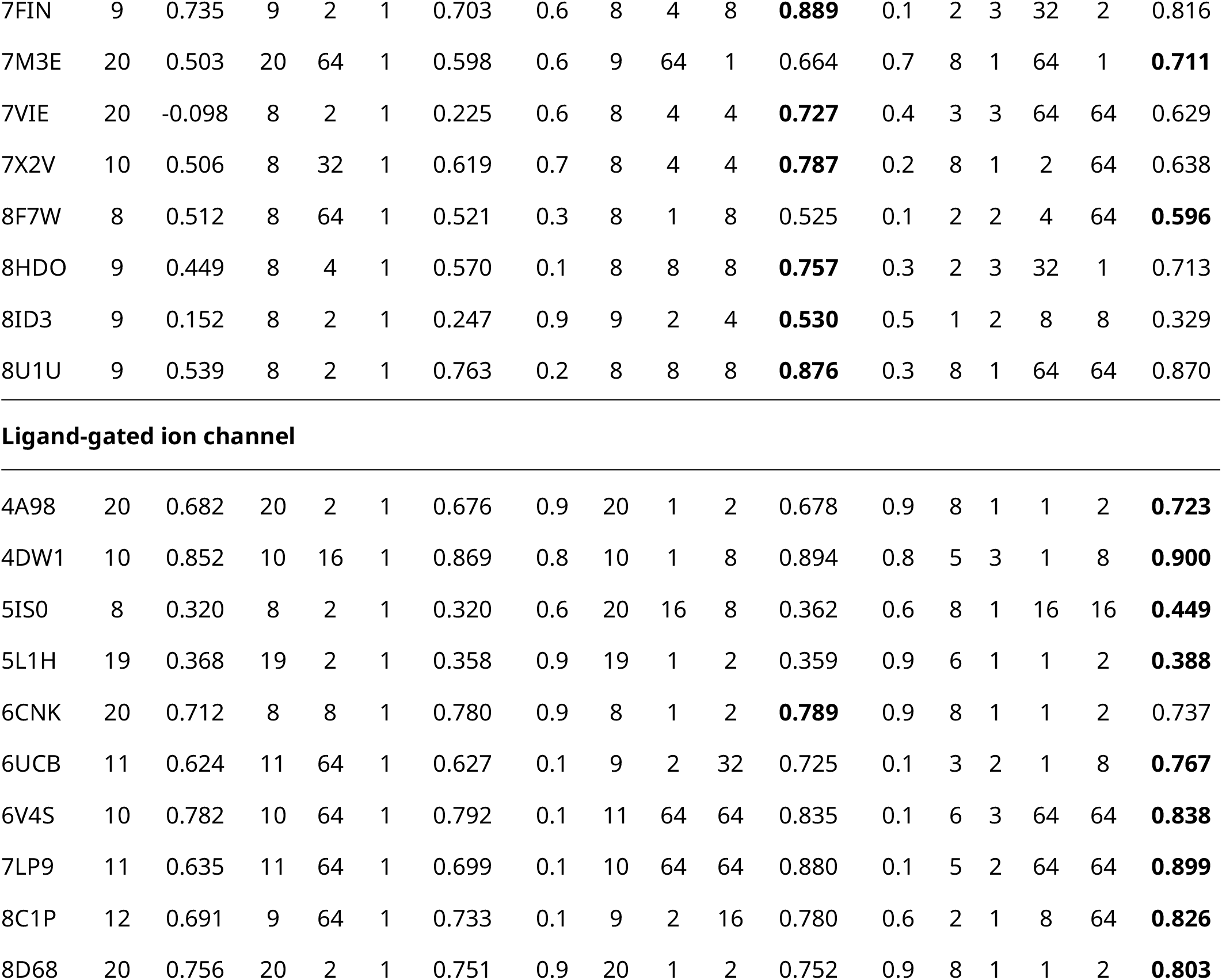
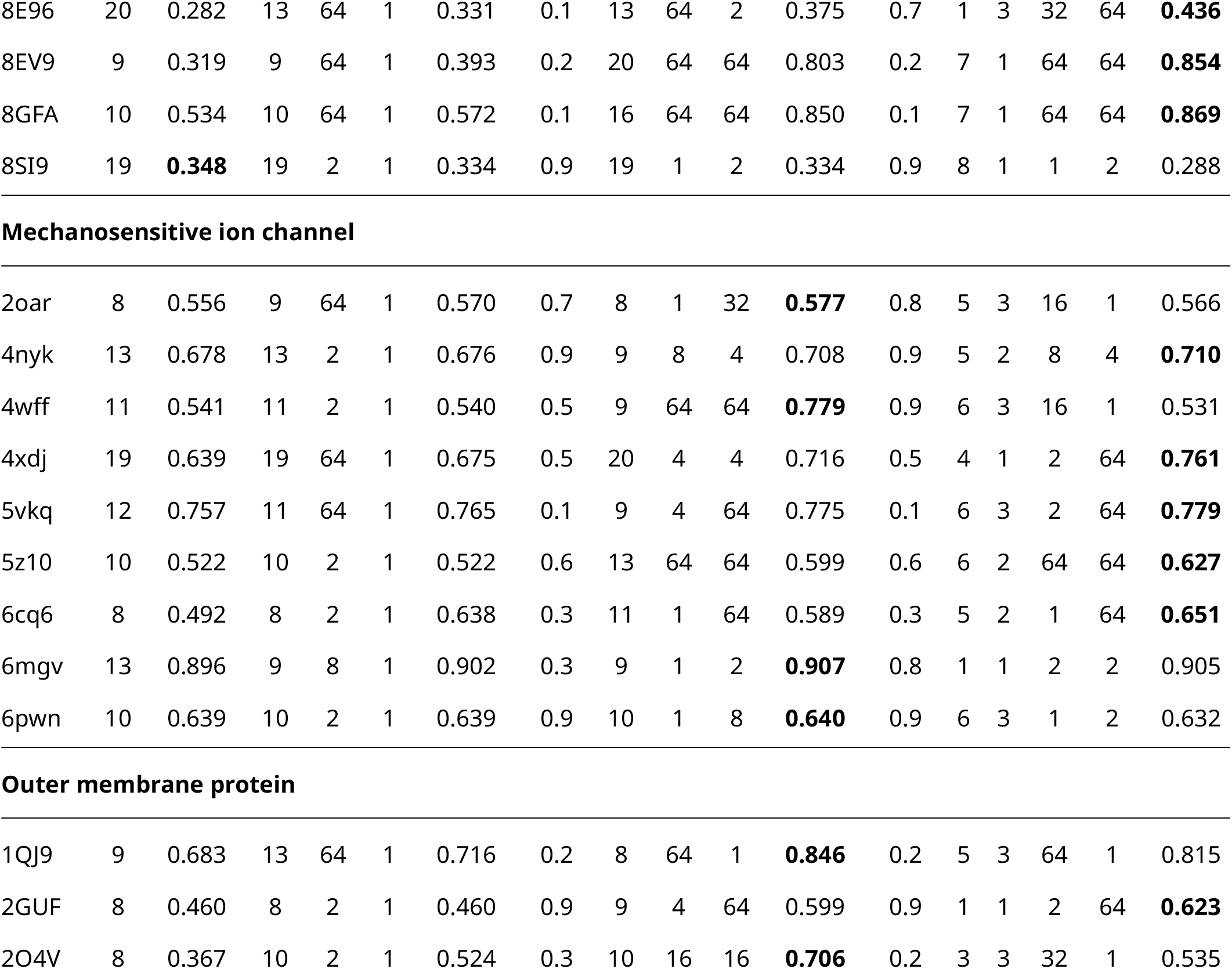
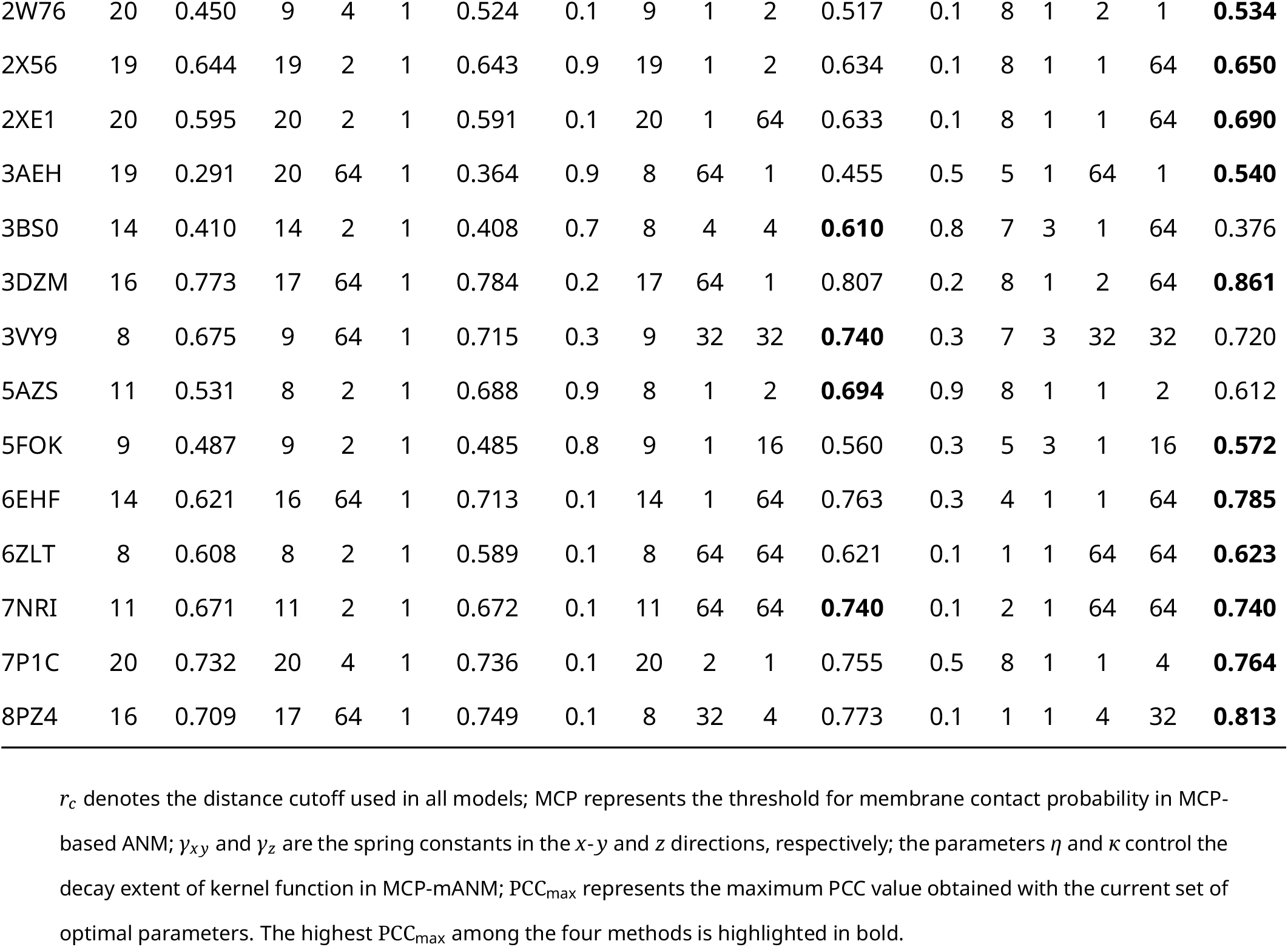
Dataset and optimized parameters.

| PDB ID | ANM |  | imANM |  |  |  | MCP-ANM |  |  |  |  | MCP-mANM |  |  |  |  |  |
| --- | --- | --- | --- | --- | --- | --- | --- | --- | --- | --- | --- | --- | --- | --- | --- | --- | --- |
| | $r_c$ | $PCC_{\max}$ | $r_c$ | $\gamma_{xy}$ | $\gamma_z$ | $PCC_{\max}$ | MCP | $r_c$ | $\gamma_{xy}$ | $\gamma_z$ | $PCC_{\max}$ | MCP | $\eta$ | $\kappa$ | $\gamma_{xy}$ | $\gamma_z$ | $PCC_{\max}$ |
| **ABC Transporter** |  |  |  |  |  |  |  |  |  |  |  |  |  |  |  |  |  |
| 4F4C | 16 | 0.410 | 16 | 64 | 1 | 0.490 | 0.9 | 16 | 32 | 2 | 0.556 | 0.4 | 3 | 1 | 64 | 1 | **0.609** |
| 4FI3 | 20 | 0.357 | 20 | 64 | 1 | 0.366 | 0.7 | 8 | 8 | 8 | **0.733** | 0.8 | 8 | 1 | 2 | 1 | 0.432 |
| 4KHZ | 20 | 0.633 | 20 | 2 | 1 | 0.634 | 0.9 | 20 | 1 | 2 | 0.629 | 0.9 | 6 | 1 | 1 | 2 | **0.656** |
| 4XTC | 20 | 0.557 | 20 | 64 | 1 | 0.592 | 0.9 | 9 | 2 | 32 | **0.653** | 0.9 | 6 | 1 | 64 | 1 | 0.633 |
| 5KC4 | 20 | 0.612 | 20 | 64 | 1 | 0.652 | 0.3 | 9 | 2 | 64 | 0.757 | 0.2 | 1 | 1 | 16 | 64 | **0.767** |
| 6QV1 | 10 | 0.632 | 8 | 2 | 1 | 0.641 | 0.1 | 8 | 64 | 64 | **0.802** | 0.9 | 8 | 1 | 1 | 2 | 0.593 |
| 6S8H | 20 | 0.672 | 9 | 2 | 1 | 0.753 | 0.9 | 20 | 64 | 1 | 0.725 | 0.9 | 8 | 1 | 4 | 1 | **0.768** |
| 6SWR | 13 | 0.748 | 13 | 8 | 1 | 0.754 | 0.9 | 8 | 1 | 16 | 0.761 | 0.9 | 3 | 2 | 8 | 64 | **0.794** |
| 6V9Z | 8 | 0.483 | 8 | 64 | 1 | 0.487 | 0.1 | 20 | 64 | 1 | 0.561 | 0.1 | 8 | 1 | 2 | 64 | **0.608** |
| 7AHC | 9 | 0.890 | 9 | 4 | 1 | 0.893 | 0.8 | 10 | 16 | 32 | 0.896 | 0.5 | 8 | 2 | 32 | 64 | **0.897** |
| 7BH2 | 10 | 0.701 | 10 | 2 | 1 | 0.698 | 0.1 | 10 | 64 | 64 | 0.757 | 0.1 | 6 | 3 | 64 | 64 | **0.778** |
| 7LKZ | 20 | 0.692 | 20 | 32 | 1 | 0.695 | 0.9 | 20 | 4 | 2 | 0.698 | 0.9 | 5 | 1 | 2 | 4 | **0.723** |
| 7O16 | 9 | 0.855 | 9 | 8 | 1 | 0.857 | 0.7 | 9 | 4 | 2 | 0.859 | 0.7 | 5 | 3 | 4 | 4 | **0.877** |
| 7P06 | 20 | 0.535 | 20 | 2 | 1 | 0.534 | 0.9 | 20 | 2 | 1 | 0.528 | 0.8 | 2 | 1 | 2 | 1 | **0.577** |
| 7R8E | 20 | 0.650 | 8 | 4 | 1 | 0.783 | 0.2 | 8 | 32 | 16 | 0.803 | 0.7 | 8 | 1 | 16 | 1 | **0.812** |
| 7VFI | 9 | 0.515 | 9 | 64 | 1 | 0.534 | 0.1 | 9 | 1 | 16 | 0.666 | 0.1 | 3 | 2 | 1 | 64 | **0.694** |
| 8FEE | 9 | 0.564 | 10 | 64 | 1 | 0.510 | 0.1 | 9 | 1 | 16 | **0.636** | 0.1 | 7 | 3 | 64 | 1 | 0.599 |
| 8G4C | 14 | 0.716 | 8 | 32 | 1 | 0.757 | 0.8 | 8 | 64 | 32 | **0.773** | 0.1 | 6 | 3 | 64 | 1 | 0.765 |
| 8HVH | 9 | 0.828 | 9 | 2 | 1 | 0.827 | 0.9 | 9 | 2 | 1 | 0.825 | 0.9 | 4 | 2 | 2 | 1 | **0.847** |
| Aquaporin |  |  |  |  |  |  |  |  |  |  |  |  |  |  |  |  |  |
| 1H6I | 9 | 0.441 | 9 | 2 | 1 | 0.441 | 0.8 | 8 | 4 | 64 | **0.505** | 0.1 | 8 | 1 | 64 | 64 | 0.466 |
| 3CN6 | 11 | 0.591 | 16 | 64 | 1 | 0.643 | 0.4 | 19 | 64 | 1 | 0.633 | 0.1 | 5 | 3 | 1 | 64 | **0.654** |
| GPCR |  |  |  |  |  |  |  |  |  |  |  |  |  |  |  |  |  |
| 4OO9 | 20 | 0.294 | 20 | 64 | 1 | 0.370 | 0.5 | 8 | 64 | 1 | 0.528 | 0.4 | 3 | 3 | 64 | 1 | **0.739** |
| 6D27 | 16 | 0.410 | 10 | 8 | 1 | 0.529 | 0.9 | 10 | 64 | 64 | **0.770** | 0.6 | 3 | 3 | 4 | 4 | 0.628 |
| 6HLP | 8 | 0.837 | 8 | 2 | 1 | 0.837 | 0.3 | 8 | 1 | 2 | **0.843** | 0.3 | 3 | 2 | 1 | 2 | 0.842 |
| 6NI3 | 9 | 0.223 | 8 | 64 | 1 | 0.371 | 0.6 | 8 | 4 | 4 | **0.832** | 0.4 | 4 | 3 | 1 | 16 | 0.510 |
| 6OIJ | 9 | 0.365 | 10 | 2 | 1 | 0.374 | 0.8 | 10 | 32 | 32 | 0.530 | 0.7 | 3 | 3 | 2 | 2 | **0.702** |
| 6X18 | 19 | 0.025 | 19 | 2 | 1 | 0.025 | 0.3 | 14 | 64 | 8 | 0.165 | 0.5 | 1 | 3 | 16 | 4 | **0.186** |
| 7AUE | 9 | 0.239 | 8 | 2 | 1 | 0.487 | 0.2 | 8 | 8 | 8 | **0.894** | 0.2 | 1 | 2 | 16 | 16 | 0.862 |
| 7C61 | 15 | -0.210 | 15 | 32 | 1 | -0.121 | 0.4 | 8 | 64 | 1 | -0.129 | 0.4 | 1 | 3 | 1 | 2 | **0.192** |
| 7F1T | 9 | 0.597 | 8 | 64 | 1 | 0.630 | 0.4 | 14 | 32 | 1 | **0.683** | 0.4 | 8 | 2 | 64 | 1 | 0.674 |
| 7FIN | 9 | 0.735 | 9 | 2 | 1 | 0.703 | 0.6 | 8 | 4 | 8 | **0.889** | 0.1 | 2 | 3 | 32 | 2 | 0.816 |
| 7M3E | 20 | 0.503 | 20 | 64 | 1 | 0.598 | 0.6 | 9 | 64 | 1 | 0.664 | 0.7 | 8 | 1 | 64 | 1 | **0.711** |
| 7VIE | 20 | -0.098 | 8 | 2 | 1 | 0.225 | 0.6 | 8 | 4 | 4 | **0.727** | 0.4 | 3 | 3 | 64 | 64 | 0.629 |
| 7X2V | 10 | 0.506 | 8 | 32 | 1 | 0.619 | 0.7 | 8 | 4 | 4 | **0.787** | 0.2 | 8 | 1 | 2 | 64 | 0.638 |
| 8F7W | 8 | 0.512 | 8 | 64 | 1 | 0.521 | 0.3 | 8 | 1 | 8 | 0.525 | 0.1 | 2 | 2 | 4 | 64 | **0.596** |
| 8HDO | 9 | 0.449 | 8 | 4 | 1 | 0.570 | 0.1 | 8 | 8 | 8 | **0.757** | 0.3 | 2 | 3 | 32 | 1 | 0.713 |
| 8ID3 | 9 | 0.152 | 8 | 2 | 1 | 0.247 | 0.9 | 9 | 2 | 4 | **0.530** | 0.5 | 1 | 2 | 8 | 8 | 0.329 |
| 8U1U | 9 | 0.539 | 8 | 2 | 1 | 0.763 | 0.2 | 8 | 8 | 8 | **0.876** | 0.3 | 8 | 1 | 64 | 64 | 0.870 |
| Ligand-gated ion channel |  |  |  |  |  |  |  |  |  |  |  |  |  |  |  |  |  |
| 4A98 | 20 | 0.682 | 20 | 2 | 1 | 0.676 | 0.9 | 20 | 1 | 2 | 0.678 | 0.9 | 8 | 1 | 1 | 2 | **0.723** |
| 4DW1 | 10 | 0.852 | 10 | 16 | 1 | 0.869 | 0.8 | 10 | 1 | 8 | 0.894 | 0.8 | 5 | 3 | 1 | 8 | **0.900** |
| 5IS0 | 8 | 0.320 | 8 | 2 | 1 | 0.320 | 0.6 | 20 | 16 | 8 | 0.362 | 0.6 | 8 | 1 | 16 | 16 | **0.449** |
| 5L1H | 19 | 0.368 | 19 | 2 | 1 | 0.358 | 0.9 | 19 | 1 | 2 | 0.359 | 0.9 | 6 | 1 | 1 | 2 | **0.388** |
| 6CNK | 20 | 0.712 | 8 | 8 | 1 | 0.780 | 0.9 | 8 | 1 | 2 | **0.789** | 0.9 | 8 | 1 | 1 | 2 | 0.737 |
| 6UCB | 11 | 0.624 | 11 | 64 | 1 | 0.627 | 0.1 | 9 | 2 | 32 | 0.725 | 0.1 | 3 | 2 | 1 | 8 | **0.767** |
| 6V4S | 10 | 0.782 | 10 | 64 | 1 | 0.792 | 0.1 | 11 | 64 | 64 | 0.835 | 0.1 | 6 | 3 | 64 | 64 | **0.838** |
| 7LP9 | 11 | 0.635 | 11 | 64 | 1 | 0.699 | 0.1 | 10 | 64 | 64 | 0.880 | 0.1 | 5 | 2 | 64 | 64 | **0.899** |
| 8C1P | 12 | 0.691 | 9 | 64 | 1 | 0.733 | 0.1 | 9 | 2 | 16 | 0.780 | 0.6 | 2 | 1 | 8 | 64 | **0.826** |
| 8D68 | 20 | 0.756 | 20 | 2 | 1 | 0.751 | 0.9 | 20 | 1 | 2 | 0.752 | 0.9 | 8 | 1 | 1 | 2 | **0.803** |
| 8E96 | 20 | 0.282 | 13 | 64 | 1 | 0.331 | 0.1 | 13 | 64 | 2 | 0.375 | 0.7 | 1 | 3 | 32 | 64 | **0.436** |
| 8EV9 | 9 | 0.319 | 9 | 64 | 1 | 0.393 | 0.2 | 20 | 64 | 64 | 0.803 | 0.2 | 7 | 1 | 64 | 64 | **0.854** |
| 8GFA | 10 | 0.534 | 10 | 64 | 1 | 0.572 | 0.1 | 16 | 64 | 64 | 0.850 | 0.1 | 7 | 1 | 64 | 64 | **0.869** |
| 8SI9 | 19 | **0.348** | 19 | 2 | 1 | 0.334 | 0.9 | 19 | 1 | 2 | 0.334 | 0.9 | 8 | 1 | 1 | 2 | 0.288 |
| 2oar | 8 | 0.556 | 9 | 64 | 1 | 0.570 | 0.7 | 8 | 1 | 32 | **0.577** | 0.8 | 5 | 3 | 16 | 1 | 0.566 |
| 4nyk | 13 | 0.678 | 13 | 2 | 1 | 0.676 | 0.9 | 9 | 8 | 4 | 0.708 | 0.9 | 5 | 2 | 8 | 4 | **0.710** |
| 4wff | 11 | 0.541 | 11 | 2 | 1 | 0.540 | 0.5 | 9 | 64 | 64 | **0.779** | 0.9 | 6 | 3 | 16 | 1 | 0.531 |
| 4xdj | 19 | 0.639 | 19 | 64 | 1 | 0.675 | 0.5 | 20 | 4 | 4 | 0.716 | 0.5 | 4 | 1 | 2 | 64 | **0.761** |
| 5vkq | 12 | 0.757 | 11 | 64 | 1 | 0.765 | 0.1 | 9 | 4 | 64 | 0.775 | 0.1 | 6 | 3 | 2 | 64 | **0.779** |
| 5z10 | 10 | 0.522 | 10 | 2 | 1 | 0.522 | 0.6 | 13 | 64 | 64 | 0.599 | 0.6 | 6 | 2 | 64 | 64 | **0.627** |
| 6cq6 | 8 | 0.492 | 8 | 2 | 1 | 0.638 | 0.3 | 11 | 1 | 64 | 0.589 | 0.3 | 5 | 2 | 1 | 64 | **0.651** |
| 6mgv | 13 | 0.896 | 9 | 8 | 1 | 0.902 | 0.3 | 9 | 1 | 2 | **0.907** | 0.8 | 1 | 1 | 2 | 2 | 0.905 |
| 6pwn | 10 | 0.639 | 10 | 2 | 1 | 0.639 | 0.9 | 10 | 1 | 8 | **0.640** | 0.9 | 6 | 3 | 1 | 2 | 0.632 |
| 1QJ9 | 9 | 0.683 | 13 | 64 | 1 | 0.716 | 0.2 | 8 | 64 | 1 | **0.846** | 0.2 | 5 | 3 | 64 | 1 | 0.815 |
| 2GUF | 8 | 0.460 | 8 | 2 | 1 | 0.460 | 0.9 | 9 | 4 | 64 | 0.599 | 0.9 | 1 | 1 | 2 | 64 | **0.623** |
| 2O4V | 8 | 0.367 | 10 | 2 | 1 | 0.524 | 0.3 | 10 | 16 | 16 | **0.706** | 0.2 | 3 | 3 | 32 | 1 | 0.535 |
| 2W76 | 20 | 0.450 | 9 | 4 | 1 | 0.524 | 0.1 | 9 | 1 | 2 | 0.517 | 0.1 | 8 | 1 | 2 | 1 | **0.534** |
| 2X56 | 19 | 0.644 | 19 | 2 | 1 | 0.643 | 0.9 | 19 | 1 | 2 | 0.634 | 0.1 | 8 | 1 | 1 | 64 | **0.650** |
| 2XE1 | 20 | 0.595 | 20 | 2 | 1 | 0.591 | 0.1 | 20 | 1 | 64 | 0.633 | 0.1 | 8 | 1 | 1 | 64 | **0.690** |
| 3AEH | 19 | 0.291 | 20 | 64 | 1 | 0.364 | 0.9 | 8 | 64 | 1 | 0.455 | 0.5 | 5 | 1 | 64 | 1 | **0.540** |
| 3BS0 | 14 | 0.410 | 14 | 2 | 1 | 0.408 | 0.7 | 8 | 4 | 4 | **0.610** | 0.8 | 7 | 3 | 1 | 64 | 0.376 |
| 3DZM | 16 | 0.773 | 17 | 64 | 1 | 0.784 | 0.2 | 17 | 64 | 1 | 0.807 | 0.2 | 8 | 1 | 2 | 64 | **0.861** |
| 3VY9 | 8 | 0.675 | 9 | 64 | 1 | 0.715 | 0.3 | 9 | 32 | 32 | **0.740** | 0.3 | 7 | 3 | 32 | 32 | 0.720 |
| 5AZS | 11 | 0.531 | 8 | 2 | 1 | 0.688 | 0.9 | 8 | 1 | 2 | **0.694** | 0.9 | 8 | 1 | 1 | 2 | 0.612 |
| 5FOK | 9 | 0.487 | 9 | 2 | 1 | 0.485 | 0.8 | 9 | 1 | 16 | 0.560 | 0.3 | 5 | 3 | 1 | 16 | **0.572** |
| 6EHF | 14 | 0.621 | 16 | 64 | 1 | 0.713 | 0.1 | 14 | 1 | 64 | 0.763 | 0.3 | 4 | 1 | 1 | 64 | **0.785** |
| 6ZLT | 8 | 0.608 | 8 | 2 | 1 | 0.589 | 0.1 | 8 | 64 | 64 | 0.621 | 0.1 | 1 | 1 | 64 | 64 | **0.623** |
| 7NRI | 11 | 0.671 | 11 | 2 | 1 | 0.672 | 0.1 | 11 | 64 | 64 | **0.740** | 0.1 | 2 | 1 | 64 | 64 | **0.740** |
| 7P1C | 20 | 0.732 | 20 | 4 | 1 | 0.736 | 0.1 | 20 | 2 | 1 | 0.755 | 0.5 | 8 | 1 | 1 | 4 | **0.764** |
| 8PZ4 | 16 | 0.709 | 17 | 64 | 1 | 0.749 | 0.1 | 8 | 32 | 4 | 0.773 | 0.1 | 1 | 1 | 4 | 32 | **0.813** |

$r_c$ denotes the distance cutoff used in all models; MCP represents the threshold for membrane contact probability in MCP-based ANM; $\gamma_{xy}$ and $\gamma_z$ are the spring constants in the x-y and z directions, respectively; the parameters $\eta$ and $\kappa$ control the decay extent of kernel function in MCP-mANM; $PCC_{\max}$ represents the maximum PCC value obtained with the current set of optimal parameters. The highest $PCC_{\max}$ among the four methods is highlighted in bold.

